# USP37 prevents unscheduled replisome unloading through MCM complex deubiquitination

**DOI:** 10.1101/2024.09.03.610997

**Authors:** Derek L. Bolhuis, Dalia Fleifel, Thomas Bonacci, Xianxi Wang, Brandon L. Mouery, Jeanette Gowen Cook, Nicholas G. Brown, Michael J. Emanuele

## Abstract

The CMG helicase (CDC45-MCM2-7-GINS) unwinds DNA as a component of eukaryotic replisomes. Replisome (dis)assembly is tightly coordinated with cell cycle progression to ensure genome stability. However, factors that prevent premature CMG unloading and replisome disassembly are poorly described. Since disassembly is catalyzed by ubiquitination, deubiquitinases (DUBs) represent attractive candidates for safeguarding against untimely and deleterious CMG unloading. We combined a targeted loss-of-function screen with quantitative, single-cell analysis to identify human USP37 as a key DUB preventing replisome disassembly. We demonstrate that USP37 maintains active replisomes on S-phase chromatin and promotes normal cell cycle progression. Proteomics and enzyme assays revealed USP37 interacts with the CMG complex to deubiquitinate MCM7, thus antagonizing replisome disassembly. Significantly, USP37 protects normal epithelial cells from oncoprotein-induced replication stress. Our findings reveal USP37 to be critical to the maintenance of replisomes in S-phase and suggest USP37-targeting as a potential strategy for treating malignancies with defective DNA replication control.

## INTRODUCTION

The eukaryotic cell cycle is a series of highly coordinated events that ensure successful genome transmission to daughter cells. DNA replication occurs during S phase of the cell cycle and is tightly regulated to achieve complete genome duplication and maintain genome integrity^1^. Accordingly, many cancers with aberrant control over cell cycle and proliferation exhibit defects in specific aspects of DNA replication. These defects often lead to replication stress and genome instability, a hallmark, driver, and exploitable target in cancer. Therefore, uncovering molecular mechanisms underlying control of DNA replication is of paramount importance to understanding fundamental aspects of cell proliferation,^2^ as well as therapeutic vulnerabilities in cancer^3^.

Broadly, DNA replication in S phase can be divided into three steps: initiation, elongation, and termination^4, 5^. Initiation occurs at specific sites known as replication origins. During G1 phase, minichromosome maintenance (MCM) complexes are loaded onto DNA throughout the genome as inactive double hexamers to ‘license’ origins for DNA replication in S phase^6^. During S-phase, the **<u>C</u>**DC45 protein associates with a subset of loaded **<u>M</u>**CM complexes along with the **<u>G</u>**INS complex (SLD5 and PLSF1-3) to generate active CMG helicases^7^. The heterodimeric TIPIN-Timeless complex is also tightly associated with the replicative helicase and contributes to both replisome and genome stability^8, 9^. As individual origins initiate replication, CMG helicases are assembled into replisomes, large macro-molecular complexes which are directly responsible for DNA unwinding and copying during S-phase^10^. During elongation, two eukaryotic replisomes unwind DNA, moving in opposite directions and synthesizing DNA.

Replication termination prior to the start of mitosis is essential for chromosome segregation fidelity and the maintenance of genome stability^11, 12^. The termination process starts when helicases either collide or reach the end of a DNA strand. Importantly, replisome unloading relies on the ubiquitin system and the action of at least two known E3 ubiquitin ligases. During a normal S-phase, unloading is controlled by the cullin RING ubiquitin Ligase CRL2^LRR1^ ^13–15^. In response to inter-strand crosslinks, unloading is coordinated by the RING E3 ligase TRAIP^15–18^. Both scenarios result in the ubiquitination of MCM7, which then recruits the conserved AAA+ ATPase enzyme p97 (also known as VCP in metazoans or Cdc48 in budding yeast)^13, 14, 19, 20^. The segregase activity of p97/VCP drives extraction and disassembly of replisomes (Fig. 1A). It is currently unknown if other signals mediate the process.

**Figure 1.**
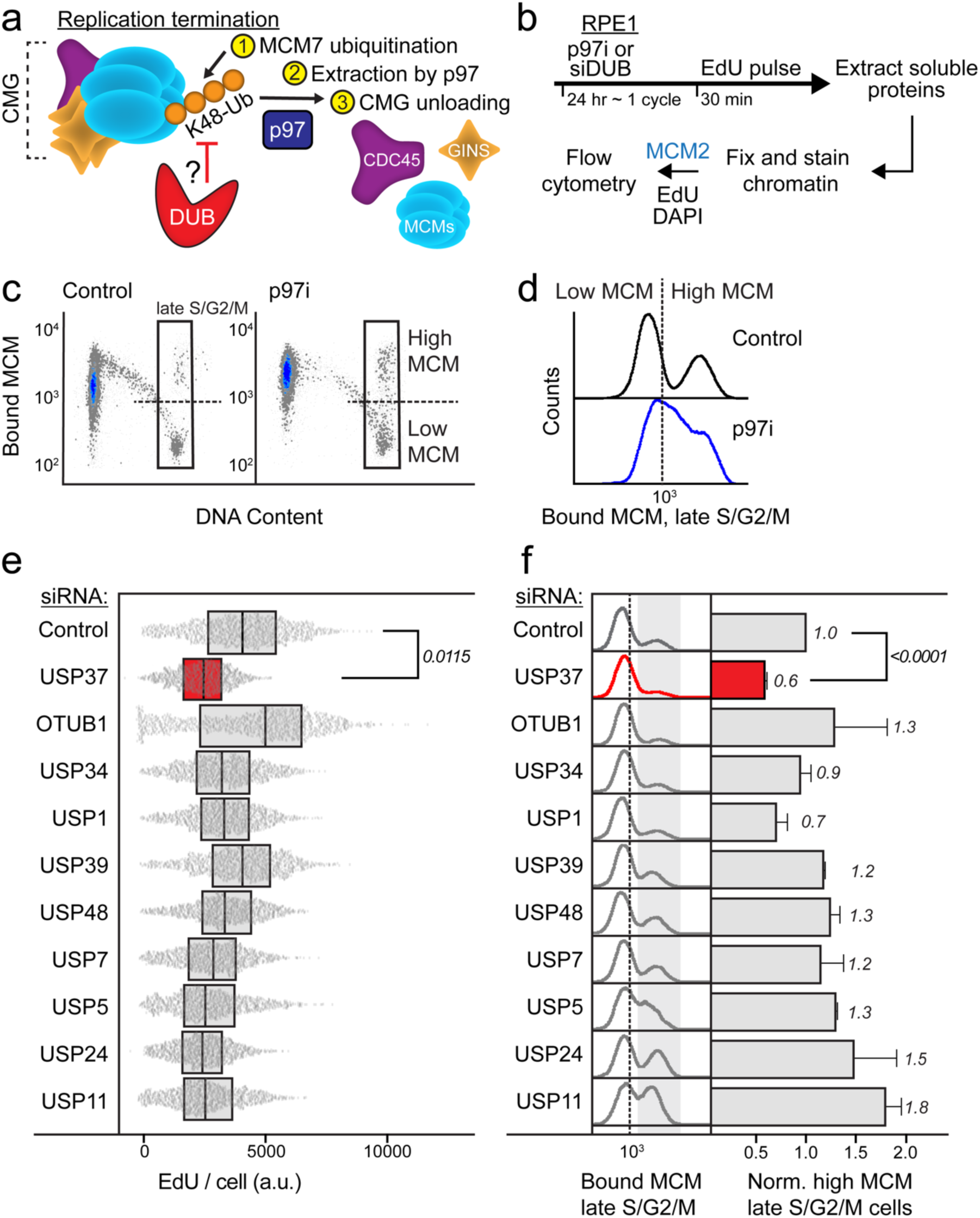
A targeted siRNA screen identifies USP37 as the antagonizing DUB for replisome disassembly. **A.** Model displaying the molecular players involved in replication termination. During replication termination, the CMG replicative helicase (CDC45-MCM2-7-GINS) is poly-ubiquitinated with lysine 48 (K48) ubiquitin linkages and the replisome is disassembled through p97. A deubiquitinase (DUB) could antagonize the ubiquitination-dependent disassembly to prevent premature replisome disassembly. **B.** Workflow for chromatin flow cytometry assays to study replication and bound MCM. RPE1-hTert cells were treated for 24 hours with either p97i or a panel of siRNAs to knock down selected DUBs individually (siDUB). Cells were labelled with EdU (thymidine analog) 30 minutes prior to harvesting, then soluble proteins were pre-extracted to retain only chromatin-bound proteins such as MCM2 (one of the replisome components). Cells were then fixed and stained for EdU (for active DNA synthesis), MCM2 (as a representative subunit for the MCM2-7 complex), and DAPI (for total DNA content) for flow cytometric analysis. **C.** Chromatin flow cytometry for RPE1-hTert cells treated with 20 nM siControl or 1.25 μM of CB-5083 (p97 inhibitor) for 24 hours, and pulsed with EdU for 30 min before harvesting. Cells were stained for bound MCM2, and DAPI (for DNA content). In the late S/G2/M gate, control cells are divided into high (>10^3^) versus low (<10^3^) bound MCM. Representative of two biological replicates. **D.** Histograms of the late S/G2/M-MCM^DNA^-positive cells from (C). **E.** RPE1-hTert cells were treated with siControl or siDUB at 20 nM as indicated. Box and whisker plots for EdU intensity per cell in S phase. Cells in each sample were randomly down sampled to 2400 cells per sample. Data is combined from two independent biological replicates. Relative fold-change of the means of EdU intensity from the two replicates was computed: siControl versus siUSP37, unpaired two tailed t-test, p=0.0115. **F.** Bound MCM in late S/G2/M from cells treated as in (E). Left: Histograms of normalized counts of the late S/G2/M-MCM^DNA^-positive cells. Right: Relative percentage of high MCM, late S/G2/M-MCM^DNA^-positive cells from at least two independent biological replicates; mean with error bars ± SEM. Unpaired two tailed t-test for the means of the three replicates for siControl versus siUSP37, p<0.0001.

As cells progress through S phase, replication forks can encounter various stressors that stall their progression. Since MCM loading is strictly prohibited during S phase to avoid re-replication, cells load excess MCM complexes onto chromatin in G1^21, 22^. A small percentage of these chromatin-bound MCM complexes are converted into active CMG helicases as part of replisomes. The excess loaded MCM serves as a reservoir of licensed “dormant origins” that can fire if cells encounter replication stress during S phase^23, 24^. Thus, it is critical to preserve both actively progressing replication forks and the reservoir of unfired origins which will be needed in the event of replication stress. Therefore, preventing the premature unloading of both replisomes and loaded but inactive MCM complexes complements mechanisms that protect acutely blocked forks to ensure faithful genome duplication^25^.

To prevent premature replication termination and maintain replication fork progression, MCM7 ubiquitination is restricted. Current models suggest that cullin-RING ubiquitin ligases (SCF^Dia2^ and CRL2^LRR1^ in budding yeast and metazoans, respectively) differentiate between actively elongating and terminated replisomes by the presence of the excluded DNA single strand^26, 27^. Interestingly, interfering with the disassembly process reduces the rate of DNA replication by impairing the recycling of replisome subunits to allow for firing of replication origins during late S-phase^28^. Moreover, because origins are only licensed in G1 and fired once and only once in S phase, there are few opportunities to recover from inappropriate replisome disassembly. We therefore postulated that previously unknown safeguards are likely to prevent premature CMG unloading and early termination.

The ubiquitination system is comprised of a trienzyme cascade (E1-E2-E3 enzymes) to target protein substrates, modulating their half-life, localization, or complex formation^29^. Deubiquitinases (DUBs) are catalytic proteases that trim or remove ubiquitin from substrates and are often critical regulators of ubiquitin signaling cascades^30^. Ubiquitination is balanced between the activity of E3 ubiquitin ligases and DUBs, with many examples of mutual regulation of targets.

The regulation of CMG unloading by ubiquitination of the MCM7 subunit suggests that DUBs could antagonize that process to prevent premature replisome unloading in S-phase. However, it is currently unknown if the ubiquitination of MCM7, and therefore replication termination, is also regulated by DUBs (Fig. 1A). Here, we identify a DUB of ubiquitin specific protease family, USP37, as a key enzyme that antagonizes MCM ubiquitination and replisome disassembly. Using a targeted loss-of-function screen, we found that USP37 stabilizes total MCM and active CMG present on chromatin. Additionally, we found that USP37 associates with the replisome machinery and restricts its disassembly through deubiquitinating MCM7. Consistent with previous data showing increased replication stress in the absence of USP37, our results indicate that USP37-mediated deubiquitination of MCM7 is essential to prevent premature replisome loss and ensure faithful DNA replication^31–34^. Because DUBs are potential therapeutic targets, we demonstrate that loss of USP37 is detrimental to cells expressing two oncoproteins that induce replication stress, cyclin E1 and c-MYC. Our findings highlight USP37 as an essential safeguard for replication fidelity and suggest a possible role in cancer pathophysiology and treatment.

## RESULTS

### A targeted siRNA screen identifies USP37 as a regulator of replisome disassembly

To identify negative regulators of MCM complex chromatin binding in S phase, we employed a previously established single-cell flow cytometry technique to quantify the amount of chromatin-bound MCM during the cell cycle (Fig. 1B)^35^. Briefly, following a pulse of the thymidine analog EdU, we used a high-salt detergent buffer to remove soluble, non-chromatin-bound proteins, leaving only chromatin bound proteins. After fixation, we detected EdU by click-chemistry to measure active DNA replication, stained cells with DAPI to determine DNA content, and immunolabeled MCM2 as a representative marker for the DNA-bound MCM2-7 complex. We analyzed non-transformed, hTERT-immortalized retinal pigment epithelial cells (RPE1) which exhibit intact cell cycle checkpoints.

To examine MCM unloading, we focused on late S and G2 phase cells which are enriched for terminating replisomes. We defined “late S/G2/M phase” cells as those with 4C DNA content and low EdU incorporation relative to mid-S phase (50% of maximum); this population also includes M phase cells. We analyzed bound MCM in this defined sub-population of late S/G2/M cells (complete gating scheme is shown in Supplementary Fig. 1). Of note, changes in MCM chromatin association in S, G2, and M phase reflects only MCM retention or unloading and not new MCM loading because all MCM loading is blocked outside of G1 phase^22^.

During late S phase, the chromatin-bound MCM complex, which is part of the CMG helicase, is unloaded during replication termination in a ubiquitin and p97-dependent mechanism^19, 20^. To validate that our flow cytometry assay accurately measures MCM unloading, we compared control cells to cells treated with a p97 small molecule inhibitor (p97i), CB-5083. Indeed, p97 inhibition led to an enrichment in chromatin-bound MCM in the late S/G2/M phase population (Fig. 1C). Plotting chromatin-bound MCM abundance in control cells as a histogram shows a bimodal distribution: a population with high levels of chromatin-bound MCM which have not yet undergone replisome disassembly (right peak), and a larger population with lower levels of chromatin bound MCM, which have undergone replisome disassembly (left peak) (Fig. 1D). Cells treated with the p97 inhibitor showed a wide continuum of chromatin-bound MCM, indicating failure to normally disassemble replisomes (Fig. 1D).

To determine if a DUB can antagonize ubiquitin-mediated replisome termination, we combined a targeted siRNA screen with our flow cytometry-based assay. We focused on a panel of DUBs that were previously identified to associate with actively replicating DNA in S phase by “isolation of proteins on nascent DNA” (iPOND)^36^. These included the ovarian tumor family deubiquitinase OTUB1 and several of the ubiquitin specific protease family, including USP1, USP5, USP7, USP11, USP24, USP34, USP37, USP39, and USP48. We treated RPE1 cells with either non-targeting siRNA or siRNA targeting each selected DUB. After approximately one complete cell division cycle in the presence of siRNA, cells were pulse-labelled with EdU, permeabilized, fixed, and analyzed for DNA synthesis and chromatin-bound MCM.

The rate of DNA replication during S phase in USP37-depleted cells was the lowest among all DUBs tested, as evidenced by substantially lower EdU incorporation per cell (Fig. 1E). These results suggest a critical role for USP37 in S phase progression. Significantly, among all DUBs tested, only USP37 knockdown resulted in a nearly unimodal distribution of chromatin-bound MCM in late S/G2/M (Fig. 1F, Supplementary Fig. 2). Thus, in USP37-depleted cells, nearly all late S/G2/M cells had undergone replisome disassembly (left peak), and the population of cells retaining high levels of chromatin-bound MCM was virtually undetectable (Fig. 1F, Supplementary Fig. 3B). USP37-depleted cells had the lowest level of chromatin-bound MCM in late S/G2/M cells, relative to all others tested (Fig. 1F). As expected, USP1 depletion also moderately impacted MCM retention in late S/G2/M due to known interactions with replication machinery^37–39^. Whereas some other DUBs also impacted EdU incorporation, USP37 was the only one whose depletion affected both EdU incorporation and MCM retention on chromatin in RPE1 cells. Consistently, U2OS cells depleted of USP37 showed significantly reduced chromatin-bound MCM levels (Supplementary Fig. 3A-C) and reduced EdU incorporation rate (Supplementary Fig. 3D), suggesting that its role in replisome disassembly and S phase progression is not cell type-specific. We therefore extended our investigation of USP37.

### USP37 prevents replisome disassembly in S-phase

Both active replisomes and licensed inactive (dormant) origins contain bound MCM complexes. To determine the role of USP37 in specifically delaying disassembly of active replisomes, we expanded our analysis to CDC45 – one of the core replisome components – which is only chromatin-bound at active replisomes (Fig. 2A)^40, 41^. We generated RPE1 cells stably expressing a doxycycline-inducible, siRNA-resistant version of full-length USP37 (siRNA resistant denoted by ^R^) (Fig. 2B). We selected single-cell clones that can express USP37 at near-endogenous or higher levels (Fig. 2C - lane 3). We then used flow cytometry to analyze the chromatin-bound levels of endogenous CDC45 in control cells and in cells treated with USP37-targeting siRNA, either with or without doxycycline induction (Fig. 2B). Depleting endogenous USP37 resulted in less chromatin-bound CDC45 during the entire S phase (Fig. 2D and E). Importantly, USP37^R^ expression rescued chromatin-bound CDC45 in S phase, indicating a direct and specific role for USP37 in preventing replisome disassembly (Fig. 2D and E). Moreover, USP37 expression rescued the reduction in EdU incorporation, indicating that USP37 is critical for normal replication in S phase (Fig. 2F). We consider it unlikely that these phenotypes reflect decreased origin firing because USP37 depletion activates rather than represses origin firing^33, 34^. Collectively, these experiments show that USP37 promotes active replisome retention.

**Figure 2.**
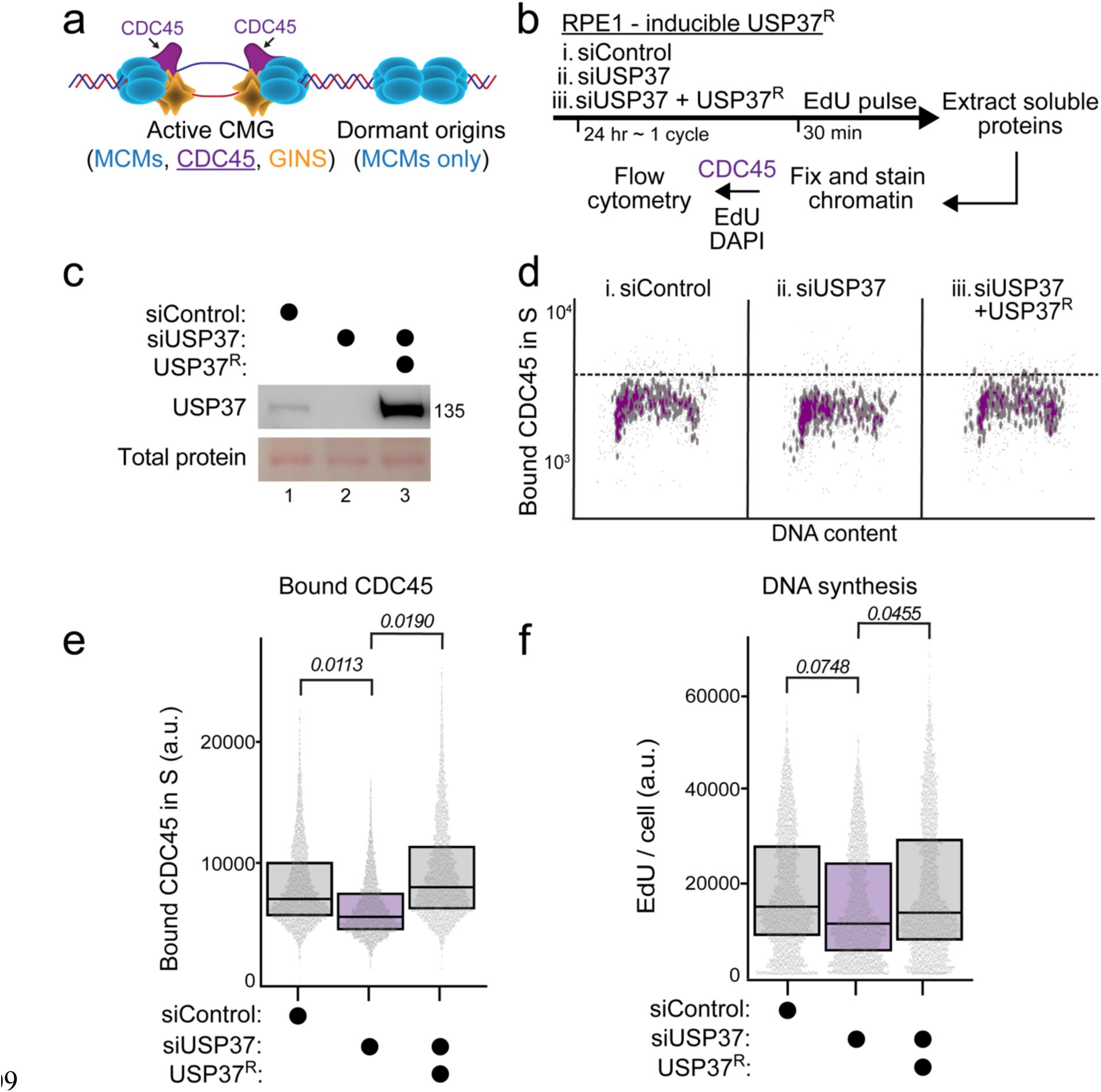
USP37 prevents replisome disassembly in S phase. **A.** Illustration of CMG at active replisomes versus MCM loaded at unfired-origins that lack CDC45. **B.** Workflow. Cells were treated with siControl or siUSP37; doxycycline was added concurrently with the siRNA treatment to express the siRNA-resistant USP37. Cells were EdU-labelled and harvested after 24 hours and analyzed by flow cytometry for endogenous bound CDC45 and DNA synthesis. **C.** Immunoblotting for endogenous and ectopic USP37 in RPE1-hTert cells treated with siControl or siUSP37 at 5 nM ± doxycycline at 20 ng/mL. Representative of four biological replicates. **D.** Chromatin flow cytometry for the same samples in (B). Cells were stained for bound CDC45, EdU incorporation (for DNA synthesis) and DAPI (for DNA content). Data shown from S phase cells only (1500 cells in each plot). Representative of four biological replicates. **E.** Quantification of (D). Box and whisker plots for chromatin-bound CDC45 intensity per cell in S phase. The aggregate of four biological replicates was randomly down sampled to ∼9600 cells per sample. Relative fold-change of the means of bound CDC45 intensity from the four replicates was computed: siControl versus siUSP37 or siUSP37 vs siUSP37 + USP37^R^, unpaired two tailed t-test, p=0.0113, 0.0190, respectively. **F.** Box and whisker plots for EdU intensity per cell in S phase from cells treated as outlined in (B). The aggregate of three biological replicates was randomly down sampled to 7500 cells per sample. Relative fold-change of the means of EdU intensity from the three replicates was computed: siControl versus siUSP37 or siUSP37 vs siUSP37 + USP37^R^, unpaired two tailed t-test, p=0.0748, 0.0455, respectively.

### USP37 interacts with replisome components

To better understand how USP37 might affect replisome disassembly, we analyzed USP37-interacting proteins by expressing FLAG-tagged USP37 (^Flag^USP37) in human embryonic kidney (HEK) 293T cells. We performed FLAG immunoprecipitation (IP) from triplicate samples followed by mass spectrometry-based proteomic analysis (Fig. 3A). Top hits from our proteomic analysis included the known USP37 interactors Cyclin A, CDH1, β-TRCP1, and β-TRCP2/FbxW11 (Fig. 3A, Supplementary table 1)^42, 43^. Remarkably, among the most enriched and statistically significant interacting proteins were all of the members of the CMG helicase complex, including MCM2-7, CDC45, GINS1-4, as well as replisome components Tipin, Timeless, and POLε (Fig. 3A and 3C). Gene Ontology (GO) analysis of the top 5% of interactors revealed strong enrichment for proteins involved in DNA replication, mitotic cell cycle progression, and the DNA damage response (Fig. 3B). We also identified additional proteins involved in DNA replication, including MCM10 and TOPBP1 (Supplementary table 1). To validate our proteomics data, we repeated our ^Flag^USP37 immunoprecipitations and immunoblotted for the replisome components CDC45, GINS1-2, MCM2, MCM7 and Timeless (Fig. 3D). Our results suggest that the USP37 deubiquitinase restricts early MCM unloading and replisome disassembly through interactions with the replisome.

**Figure 3.**
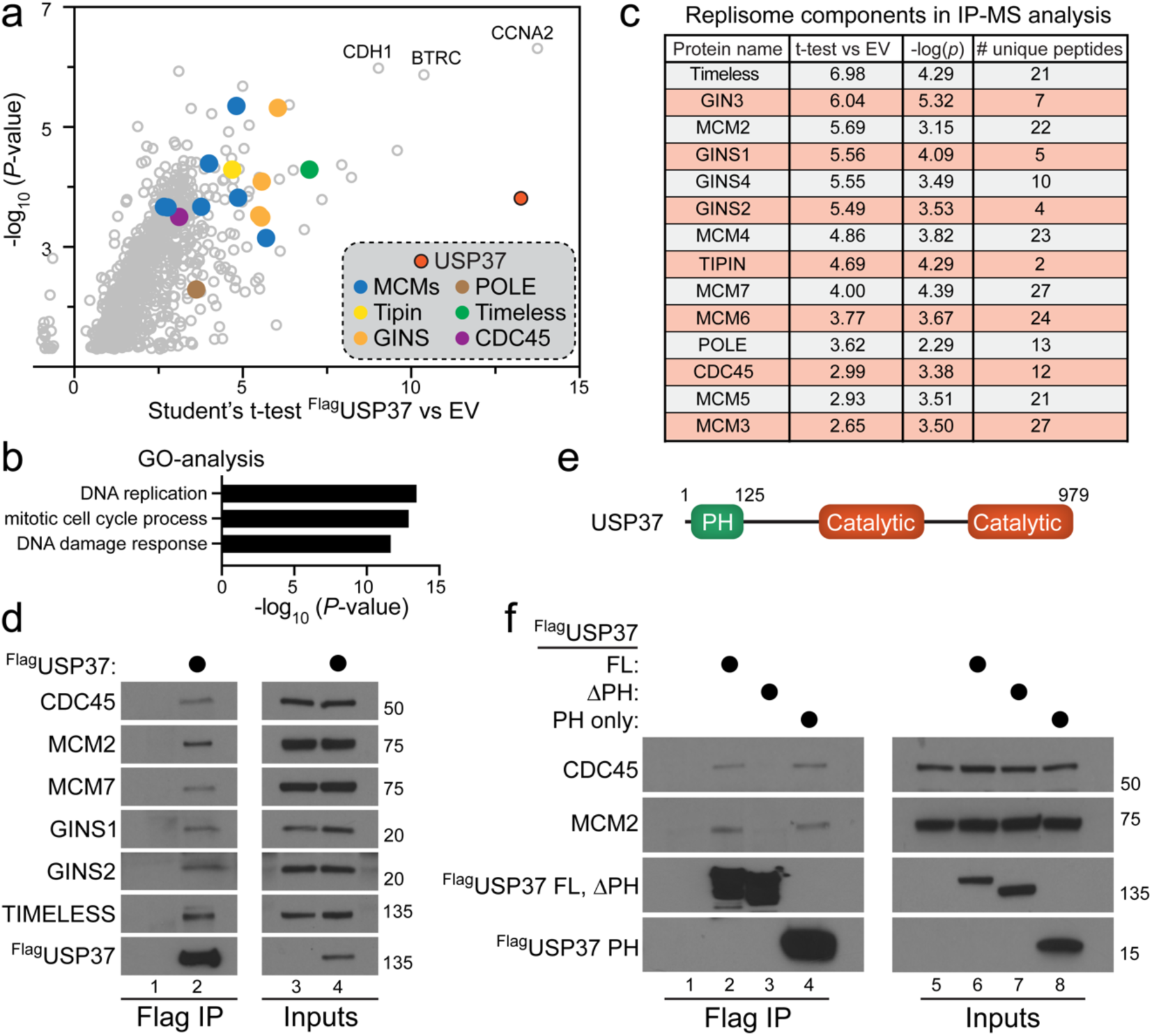
USP37 interacts with the replisome components. A. Triplicate samples of HEK293T cells expressing FLAG-tagged USP37 or an empty vector (EV) control were subjected was FLAG immunoprecipitation. Immunoprecipitates were washed, eluted and subjected to proteomic analysis. Proteins enriched in USP37 IPs, compared to controls, are shown. B. Gene Ontology analysis was performed on significantly enriched proteins to reveal the majority of USP37 interactors are involved in DNA replication and cell cycle progression. C. All components of the CMG complex were enriched in the FLAG-USP37 IP-MS. The fold enrichment over control, p-value, and number of peptides identified are shown. D. HEK293T cells were transfected for 48 hours with FLAG-tagged USP37 or an empty vector as a control. FLAG-USP37 was subjected to FLAG immunoprecipitation and analyzed by immunoblot. The indicated endogenous components of the CMG complex were co-precipitated by USP37. E. USP37 schematic. FL in (F) corresponds to full length USP37, Δ corresponds to USP37 lacking the PH domain, and PH corresponds to a USP37 fragment containing the PH domain only. F. The interaction between USP37 FL, Δ, or PH was assessed as described in D. USP37 interacts with the CMG complex through its PH domain.

The USP family of deubiquitinases is characterized by extensions and insertions into their conserved catalytic domains^44^. USP37 contains an insertion into its catalytic domain that contains three ubiquitin-interacting motifs (UIMs), important for its full catalytic activity^45, 46^. Interestingly, USP37 also contains an N-terminal extension that includes a Pleckstrin Homology (PH) domain of unknown function (Fig. 3E). The CRL2 substrate receptor responsible for MCM7 ubiquitination, LRR1, also contains an N-terminal PH domain which is required to recruit CRL2^LRR1^ to CMG, leading to MCM7 ubiquitination and replisome disassembly^26^. We therefore tested whether the USP37 PH domain is similarly important for USP37 binding to CMG. We examined whether USP37 mutants lacking the PH domain can still interact with CMG components. Importantly, a FLAG-tagged, truncated version of USP37 lacking the PH domain (ΔPH) was unable to bind CDC45, MCM2 and MCM7, whereas full-length (FL) USP37 readily bound them (Fig. 3F). Further, the PH domain alone (PH-USP37) was sufficient to bind these core replisome proteins (Fig. 3F). Expression of eGFP-tagged versions of FL-USP37 and

ΔPH-USP37 in U2OS cells showed that both localize to the nucleus (Supplementary Fig 4A). Further, the truncated version of USP37 lacking its PH domain retained its full enzymatic activity, based on its ability to react with a ubiquitin vinyl-sulfone activity-based probe (Supplementary Fig. 4B)^45^. Therefore, we conclude that the PH domain is dispensable for USP37’s catalytic activity and localization to the nucleus but is vital for the USP37-replisome interaction.

### USP37 regulates the CMG complex by deubiquitinating MCM7

It has been shown previously that replisome disassembly is triggered by MCM7 ubiquitination. Since our results indicate that USP37 preserves active replisome assembly by binding to the CMG complex, we hypothesized that MCM7 could be subjected to USP37-mediated deubiquitination. To explore this hypothesis, we first established that FLAG-tagged USP37 can interact with V5 epitope-tagged MCM7 when expressed in HEK293T cells (Fig. 4A). Next, we tested if USP37 can regulate endogenous MCM7 ubiquitination. We used an RPE1 cell line which stably expresses 6xHis-FLAG-tagged ubiquitin, allowing us to isolate ubiquitinated proteins on Ni-NTA resin under strong denaturing conditions. As shown in Figure 4A, MCM7-V5 has an apparent molecular weight of 75 kDa (input panel, lane 1). Following Ni^2+^ pull down, we observed a single slower-migrating form of endogenous MCM7 at ∼100 kDa by SDS-PAGE which corresponds to ubiquitinated MCM7 because it is absent from cells that do not express 6xHis-FLAG-Ub (Fig. 4B, compare lanes 1 and 3). Importantly, depleting USP37 using siRNA led to the appearance of additional, higher molecular weight, ubiquitinated forms of MCM7 (Fig. 4B), 6xHis-Ub pulldown panel, lane 2). Since ubiquitinated proteins were isolated under denaturing conditions, these bands represent MCM7 protein that is covalently conjugated to ubiquitin. These results suggest that under normal growth conditions, MCM7 ubiquitination is modulated by USP37.

**Figure 4.**
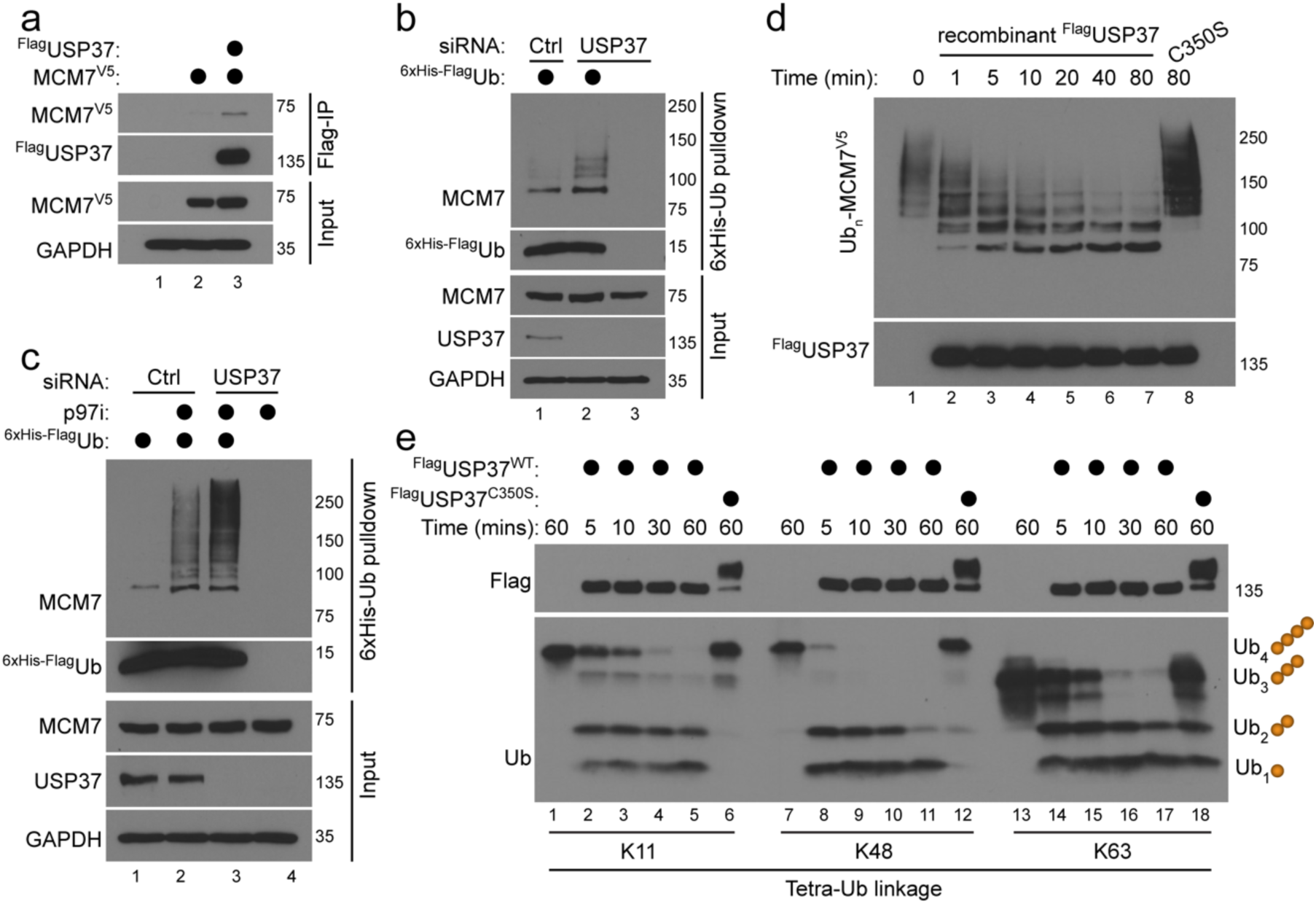
USP37 regulates the CMG complex by deubiquitinating MCM7. **A.** HEK293T cells were transfected for 48 hours with MCM7-V5, alone or in combination with FLAG-USP37. MCM7 interacts with USP37, as observed by immunoblot using the indicated antibodies. **B.** USP37 was knocked down using siRNA for 48 hours in RPE1 cells stably expressing a 6His-FLAG-tagged ubiquitin construct. Ubiquitinated proteins were pulled down using Ni-NTA, revealing that USP37 siRNA increases endogenous MCM7 ubiquitination, as observed by immunoblotting. **C.** MCM7 ubiquitination was analyzed as described in B, except that cells were treated with 5 µM of the p97i CB-5083 for the last 4 hours before harvesting. Inhibition of p97 strongly increases MCM7 ubiquitination, and this is even more pronounced after USP37 knock down. **D.** Ubiquitinated MCM7 isolated from HEK293T cells was mixed with 100 nM of recombinant USP37 WT or a catalytically inactive mutant (C350S). The in vitro deubiquitination assay shows that USP37 WT, but not C350S, deubiquitinates Ub-MCM7. **E.** Flag-tagged USP37 was ectopically expressed for 48 hours and subsequently purified from HEK-293T cells by FLAG immunoprecipitation. USP37 immunoprecipitates were mixed with 1 µM of K11, K48 or K63 tetra-ubiquitin chains, revealing that USP37 cleaves Tetra- and Tri-Ub more efficiently than Di-Ub.

It has been shown that p97 targets ubiquitinated MCM7 for replisome disassembly^19, 20^. Therefore, we tested the importance of USP37 in regulating MCM7 ubiquitination when p97 activity is inhibited. Consistent with prior reports, MCM7 ubiquitination was strongly increased in RPE1 cells treated with the p97i CB-5083 (Fig. 4C, 6xHis-Ub pulldown panel, compare lanes 1 and 2). Moreover, MCM7 ubiquitination was further increased by depleting USP37 in the presence of the p97i (Fig. 4C, lane 3). This additive effect of p97 inhibition and USP37 depletion on MCM7 ubiquitination suggests that USP37 and p97 likely function at separate steps in the process (Fig. 4C). Since p97 acts at the final step of replisome disassembly, these data collectively imply that USP37 controls unloading at a step prior to disassembly through promoting MCM7 deubiquitination.

The experiments above show that USP37 regulates MCM7 ubiquitination in cells. We next determined if the deubiquitinating activity of USP37 toward MCM7 could be recapitulated *in vitro*. Over the course of this study, we noticed that co-expression of our 6xHis-FLAG-Ubiquitin plasmid along with the V5-tagged MCM7 (MCM7-V5) in HEK-293T cells led to significant MCM7 ubiquitination, as observed by immunoblot following Ub pulldown (Supplementary Fig. 4C). Thus, we devised a strategy to isolate and elute ubiquitinated proteins, including ubiquitinated MCM7, from HEK-293T cells (see Methods). We confirmed that we recovered substantial ubiquitinated MCM7 with this approach by immunoblotting with V5 and MCM7 antibodies (Supplementary Fig. 4C). In parallel, we produced WT and catalytically inactive (C350S) versions of recombinant USP37 from baculovirus-infected insect cells (Supplementary Fig. 4D).

To test if USP37 can deubiquitinate MCM7 *in vitro*, we mixed recombinant USP37 with our ubiquitinated protein eluate and assessed MCM7 deubiquitination over time by MCM7 immunoblotting. Significantly, USP37 efficiently removed ubiquitin from MCM7. This was dependent on USP37 activity, since USP37 harboring a C-S mutation of its active site cysteine at position 350 was unable to promote MCM7 deubiquitination (Fig. 4D compare lanes 7 and 8). Thus, USP37 can deubiquitinate MCM7, establishing that MCM7 ubiquitination is directly antagonized by USP37. Interestingly, USP37 efficiently removed the high molecular weight species of ubiquitinated MCM7 (Fig. 4D, 150 kDa and above), but at the end of the reaction, MCM7 was still modified and migrated at a position consistent with the retention of ∼2-3 ubiquitin molecules (Fig. 4D, molecular weight species between 75 and 100 kDa). This observation, combined with the fact that cells treated with p97i and USP37 siRNAs showed a strong increase of high molecular weight MCM7-Ub (150 kDa and above), suggested that USP37 might prefer to deubiquitinate longer Ub chains.

To test this possibility, we isolated FLAG-USP37 from HEK293T cells and incubated it with isolated Ub chains that are 4 ubiquitins in length (tetra-Ub, “Ub_4_”). To test the linkage specificity of USP37 in this system, three Ub chain types were independently examined, containing linkages through either Lysine 11 (K11), K48, or K63. During a time course, USP37 rapidly cleaved tetra- and tri-Ub to di-Ub (Fig. 4E lanes 2-5, 8-11, and 14-17). However, USP37 hydrolyzed di-Ub to mono-Ub much more slowly (Fig. 4E). We also observed that USP37 prefers K48-linked ubiquitin chains over K11- or K63-linked ubiquitin chains (Fig. 4E). The catalytically inactive USP37-C350S mutant had no activity in these assays (Fig. 4E lanes 6, 12, and 18). This preference is significant because CRL2^LRR1^ predominately creates K48-linked Ub chains on MCM7^19,20^. Collectively, these experiments demonstrate that USP37 can deubiquitinate MCM7 and suggest that it does so by preferentially removing long ubiquitin polymers.

### USP37 protects replisomes during oncogene-driven replication stress

We analyzed coessentiality data from the cancer dependency map (DepMap) project to determine genes and pathways related to USP37 function. Consistent with our experimental data showing a role for USP37 in DNA replication, using expression-corrected CERES scores, dozens of genes involved in DNA replication strongly correlated with USP37, including numerous MCMs, CDC45, and other components of active replisomes (Fig 5A).

**Figure 5.**
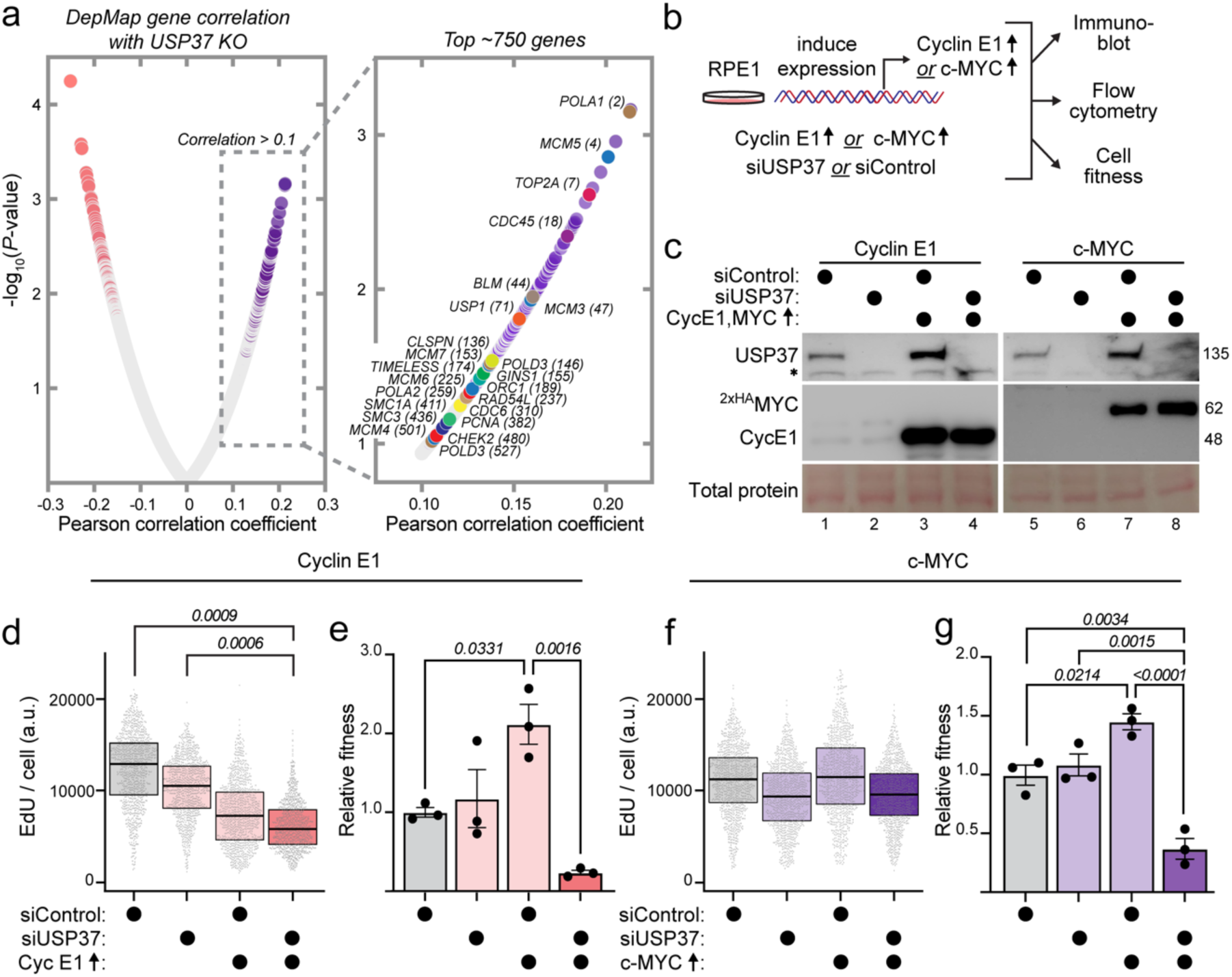
USP37 protects replication efficiency and proliferation in oncoprotein-expressing cells. **A.** Expression-corrected CERES correlation scores were downloaded from The DepMap database for genes similar to USP37 knockout. Proteins involved in DNA replication and the DNA damage response are significantly enriched. Proteins are color coded similarly to Figure 3A (MCMs = blue, CDC45 = purple, polymerases = brown). **B.** RPE1-hTert cells engineered for either doxycycline-inducible Cyclin E1 or c-MYC were treated to induce expression simultaneously with USP37 depletion to examine effects on DNA replication and cellular fitness. **C.** Immunoblotting for USP37, Cyclin E1 or c-MYC in RPE1-hTert cells as outlined in B. siControl or siUSP37 were used at 5 nM. Doxycycline was added simultaneously with the siRNA at 100 or 25 ng/mL to overproduce Cyclin E1 or c-MYC for 24h, respectively as indicated. Representative of two biological replicates. **D.** EdU intensity per cell for the experiment described in (B) to overproduce Cyclin E1. The aggregate of two biological replicates was randomly down sampled to 2000 cells per sample. Relative fold-change of the means of EdU intensity from the two replicates: siControl versus siUSP37+ Cyclin E1 or siUSP37 versus siUSP37+ Cyclin E1 or Cyclin E1 versus siUSP37+ Cyclin E1, was computed, unpaired two tailed t-test, p=0.0009, 0.0006, 0.3059, respectively. **E.** Normalized fitness for RPE1-hTert Cyclin E1 cells treated with siControl or siUSP37 with or without overexpression of Cyclin E1 to induce replication stress for five days total. n = 3 biological replicates. * = *p* ≤ 0.05, ** = *p* ≤ 0.01 by one-way ANOVA. **F.** EdU intensity per cell for the experiment described in (B) to overproduce c-MYC. The aggregate of two biological replicates was randomly down sampled to 2000 cells per sample. Relative fold-change of the means of EdU intensity from the two replicates: siControl versus siUSP37+ c-MYC or siUSP37 versus siUSP37+ c-MYC or c-MYC versus siUSP37 + c-MYC, was computed, unpaired two tailed t-test, p=0.2044, 0.7278, 0.4, respectively. **G.** Normalized fitness for RPE1-hTert c-Myc cells treated with siCTRL or siUSP37 with or without overexpression of c-MYC to induce replication stress for three days total. n = 3 biological replicates. * = *p* ≤ 0.05, ** = *p* ≤ 0.01, **** = *p* ≤ 0.001 by one-way ANOVA.

Genome instability is a hallmark of cancer. Loss of function in key genes involved in genome maintenance (e.g., *TP53* and *BRCA1)* contributes to a significant portion of cancers^47, 48^. Conversely, many oncoproteins, including cyclin E and c-MYC, induce replication stress, which generally increases reliance on the cellular pool of loaded, but unfired MCM-complexes^49^. Thus, inactivation of proteins that maintain MCM on chromatin could represent a vulnerability in cells with oncoprotein activation, suggesting that cancers undergoing replication stress could be dependent on USP37.

We hypothesized that cells with elevated replication stress would be vulnerable to loss of USP37. To test the effects of USP37 loss in the context of cyclin E or c-MYC overproduction, we generated RPE1 cells that stably express either doxycycline-inducible *CCNE1* (the gene for Cyclin E1) or c-*MYC* (Fig 5B). Upon addition of doxycycline, these proteins are overproduced (Fig 5C). Cyclin E overproduction alone reduced the rate of DNA synthesis (EdU/cell) as expected based on previously-documented defects in origin licensing^35, 50^ (Fig. 5D), and concurrently depleting USP37 caused a further decrease (Fig. 5D). Overproducing c-MYC did not substantially impact the rate of DNA synthesis in a parallel experiment (Fig. 5F). Although both cyclin E1 and c-MYC drive S phase when over-produced, their mechanisms only partially overlap which may explain why cyclin E1 impaired the rate of DNA synthesis whereas c-MYC did not (Fig. 5D and 5F) ^51–54^.On the other hand, Cyclin E and c-MYC both stimulated cell population increase as measured by an assay for total viable cell population (Fig. 5E and G). Depleting USP37 by siRNA in cyclin E-overproducing cells dramatically reduced total cell fitness (Fig 5E). Similarly, depleting USP37 in combination with c-MYC overproduction significantly reduced cell fitness (Fig 5G). Taken together, our results indicate that USP37-mediated replisome preservation is important for accommodating oncogene-induced replication stress.

## DISCUSSION

Chromosome duplication must occur completely and with high precision to prevent genome instability, a hallmark of cancer. Critical to this process is the accurate and timely assembly and disassembly of replisomes. Therefore, both loading and unloading of the CMG helicase is tightly regulated through many different signaling pathways and processes^55^. Unloading of the CMG complex during replication termination is accomplished by the CRL2^LRR1^-mediated ubiquitination of MCM7, and subsequent dismantling of the replisome by the p97 segregase^6, 11, 13, 14, 19, 20, 56^. The RING E3 TRAIP can also ubiquitinate MCM7 and does so in response to interstrand crosslinks^57^. Premature ubiquitination is prevented, in part, by the excluded DNA strand during replication^26^. However, if the LRR1-MCM interface is exposed due to any mispositioning of the excluded DNA strand, premature replisome disassembly could still occur. Thus, we hypothesized that an additional safeguard mechanism would exist to prevent premature replication termination and could be controlled at the level of MCM7 ubiquitination. Here, we demonstrate that premature CMG unloading is prevented by the USP37 deubiquitinase, which antagonizes MCM7 ubiquitination, thereby limiting aberrant unloading of the CMG helicase. Prior to our study, an enzyme that directly antagonizes MCM7 ubiquitination had not been identified.

Since the family of ∼100 DUBs enzymatically remove ubiquitin marks from substrates, we tested if a DUB protects the replisome from premature unloading. USP37 has previously been linked to S-phase and located at replication forks^32, 34^. Furthermore, numerous previous studies identified its importance in response to DNA damage and replication stress^33, 34, 58, 59^. However, a complete understanding of its role in DNA replication was not well established. Through a series of complementary approaches, we discovered that USP37 binds to the replisome, controls the level of polyubiquitin on MCM7 in cells, and prevents premature CMG unloading. We further identified the USP37 PH domain as a potential mediator of the USP37-replisome interaction. Detailed dissection of how USP37 binds to the replisome will be important to understand how the activities of LRR1 and USP37 are properly balanced. For example, as both enzymes have PH domains, it is possible that they compete for binding to the CMG.

Likewise, understanding the switch between protection by USP37 and LRR1 activity will be pivotal. How is USP37 activity quenched at or removed from CMG, how is LRR1 activated, and what are the mediating factors governing the timing of these interactions? USP37 is degraded in G2-phase, in a PLK1 and βTRCP dependent manner^43^. Thus, one possibility is that the switch from replisome stabilization to its disassembly is coordinated with cell cycle progression through the inactivation of S-phase kinases, like CHK1, whose activation is promoted by the presence of single stranded DNA and is enhanced by USP37^33, 34^. Subsequent activation of PLK1 to stimulate USP37 degradation likely accelerates ubiquitin-dependent replisome disassembly in G2.

Ubiquitin signals generated by E3 ubiquitin ligases are always potentially subject to editing by DUBs. Deubiquitination can prevent protein degradation by disassembling proteolytic signals and can also extinguish non-degradative ubiquitin signaling events. We reasoned that since ubiquitination of MCM7 triggers replisome unloading, that DUBs could antagonize complex disassembly. We show here that MCM7 ubiquitination is antagonized by USP37. Interestingly, E3 ligases can assemble ubiquitin chains linked through each of the lysines in ubiquitin (e.g., K48, K63, etc.), and can also generate branched chains. Furthermore, DUBs can edit or sculpt these ubiquitin chains or prevent the formation of branched or mixed chains. Interestingly, in our hands, USP37 reduced but did not completely abolish ubiquitination of MCM7, suggesting that it may be removing a specific chain or chain type(s), rather than deubiquitinating the proximal ubiquitin conjugated directly onto MCM7 (Fig. 4D). Therefore, it remains an open question if the ubiquitin chains formed on the replisome are simply trimmed by USP37, lowering the total amount of polyubiquitination, and/or are edited by USP37 to potentially facilitate building a qualitatively different ubiquitin signal, thereby indicating the presence of replication stress or eliciting a cellular response.

Replication stress can arise through endogenous and exogenous sources. A potential source of replication stress is re-replication. One mechanism that prevents toxic re-replication is the strict prevention of MCM loading after the start of S phase, ensuring that MCM loading occurs only in G1. However, since the total amount of loaded MCM cannot be increased once S-phase begins, cells load more MCM than is needed, and only a fraction of the total loaded MCM is converted into active CMGs in an unperturbed S-phase. Failing to load an excess of MCM complexes can be detrimental. Similarly, premature CMG unloading during S phase, prior to the completion of DNA replication, might also be detrimental.

The need for excess MCM is particularly important when cells encounter replicative stress, which can convert the backup, excess MCM into active replisomes. Replicative stress can occur in response to both chemical stressors that impact the DNA (e.g., topoisomerase inhibitors) and the activation of oncoproteins, like cyclin E1 and c-MYC. The latter induces replication stress through various mechanisms, including mismanaging CMG assembly or activation. Cyclin E1 drives premature S phase entry before enough MCMs are loaded onto chromatin in G1 phase, which results in underlicensing^50, 60^. Thus, cells overproducing cyclin E1 are depleted of the normally available backup pool of licensed origins. Moreover, it has been reported that both cyclin E and c-MYC deregulation induce premature and disrupted origin firing in intragenic regions^53, 61^. In addition, cells overproducing c-MYC over activate their licensed origins in S phase through promoting CDC45 and GINS recruitment to MCM hexamers^62^.

Excessive origin activation in S phase also depletes cells of their dormant origin pool^63^. Notably, G1 phase is the only window available for cells to license origins, and there are multiple regulatory pathways that strictly inhibit licensing in S phase to prevent re-replication^64^. Thus, in both cases of oncogene-induced CMG mismanagement, the fired origins in S phase must be protected from premature disassembly to prevent under replication and genome stability. Consistent with this idea, we demonstrate that USP37 depletion in cells overproducing cyclin E1 or c-MYC dramatically undermines S phase progression and cell fitness. We suggest this USP37-mediated mechanism complements other replication fork protection mechanisms to preserve replication capacity in S phase^65^.

From a therapeutic perspective, DNA damaging agents are often used in cancer chemotherapies. However, many have unwanted side effects. Previous reports show that USP37 depletion can sensitize cells to these agents^34, 58, 66^. This sensitization suggests that inhibiting USP37 could be advantageous, in combination with cancer therapeutics, to selectively target malignant cells. Currently, few small molecules that are potent and selective DUB inhibitors exist but identifying them remains a promising and underutilized approach for the treatment of cancer.

## ACKNOWLEDGEMENTS

We thank lab members and colleagues at UNC for helpful discussions throughout this project and the Labib, Walter, and Jackson labs for sharing results prior to publication. Our funding is as follows: M.J.E (UNC University Cancer Research Fund (UCRF); NIH R01GM134231 and R35GM153250; ACS RSG-18-220-01-TBG), N.G.B (NIH R35GM128855 and UCRF), J.G.C (NIH R35GM141833), D.L.B (NIH T32GM008570), and D.F (AHA 23PRE1027147). The UNC Flow Cytometry Core Facility (RRID:SCR_019170) is supported in part by P30 CA016086 Cancer Center Core Support Grant to the UNC Lineberger Comprehensive Cancer Center. The UNC Proteomics Core Facility is supported in part by NCI Center Core Support Grant (2P30CA016086-45) to the UNC Lineberger Comprehensive Cancer Center. Research reported in this publication was supported in part by the North Carolina Biotech Center Institutional Support Grant 2017-IDG-1025 and by the National Institutes of Health 1UM2AI30836-01.

## AUTHOR CONTRIBUTIONS

DB, DF, TB, JC, NB, and ME conceived experiments.

DB, DF, and TB carried out most cell biological and biochemical experiments.

XW performed USP37 IP for proteomic analysis.

BM generated RPE1 cells line expressing 6HIS-tagged ubiquitin.

DB, DF, TB, JC, NB, and ME contributed to the writing of the manuscript.

All authors provided input on and approved of the manuscript. …

## DATA AVAILABILITY

Proteomics data, including raw files and search parameters, were uploaded to ProteomeXchange via PRIDE (Identifier XXXX) and are available publicly.

---

**Supplementary fig. 1.**
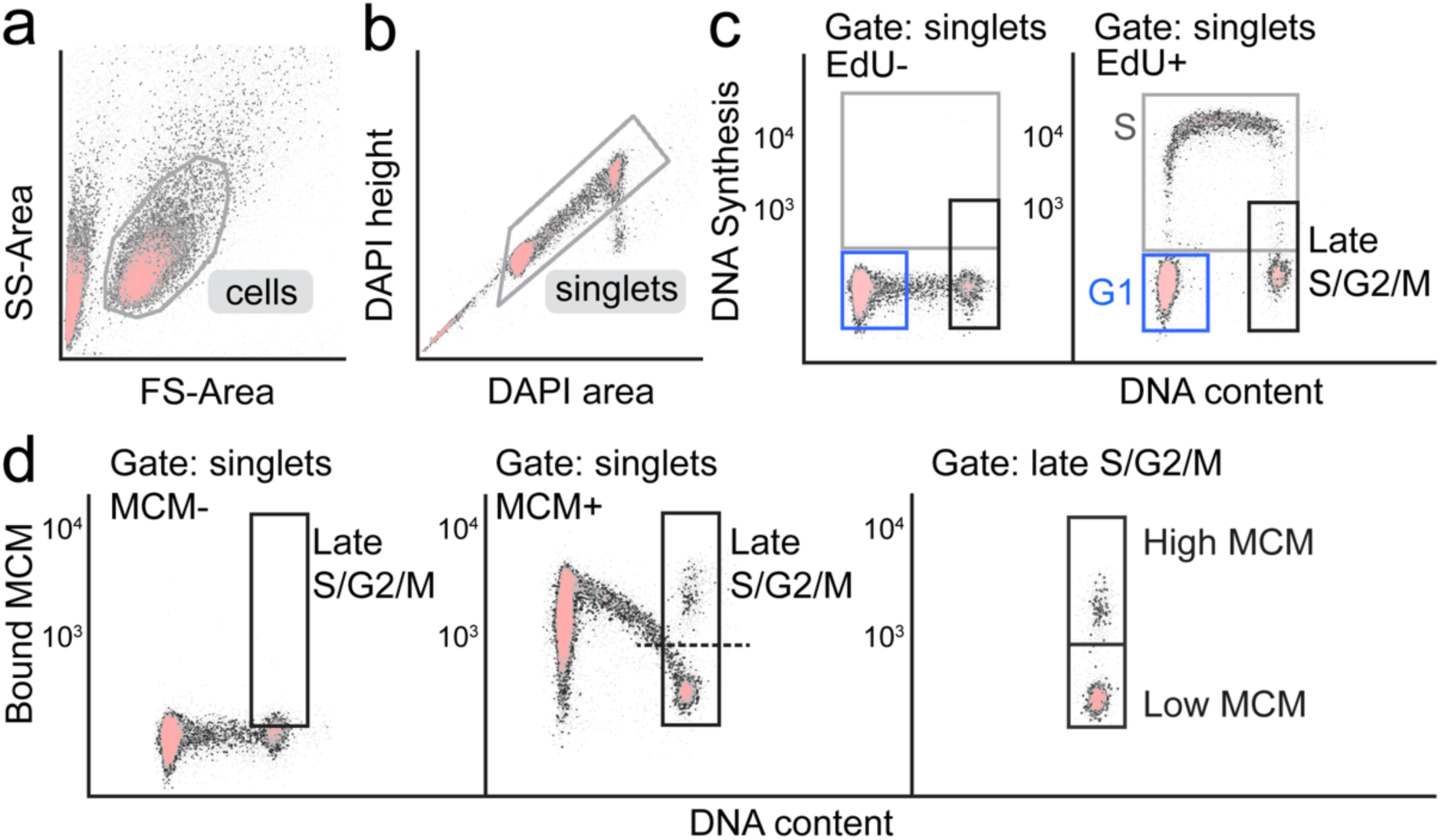
Flow cytometry gating scheme. **A.** Example of control RPE1-hTert cells. Cells are gated on FS-area versus SS-area to exclude debris. **B.** Singlets or individual cells are gated on DAPI area versus DAPI height to exclude doublets. **C.** Cell cycle phases are determined based on DAPI and EdU staining for DNA content and DNA synthesis, respectively. An EdU negative sample was used to determine the gate for S phase cells (EdU positive). G1 cells have 2C DNA content and are EdU negative. Late S/G2/M cells have 4C DNA content and EdU intensity: max: ∼50% of the max EdU intensity, and min: EdU negative. **D.** MCM2 was detected using anti-MCM2 antibody in cells stained with DAPI. Left: A negative control sample (unstained with MCM2 antibody but stained with secondary antibody and DAPI) was used to define background MCM2 staining. Middle: Chromatin extracted RPE1-hTert cells show a distribution of bound MCM during G1 phase, and a gradual decrease in bound MCM during S phase. Right: The late S/G2/M gate determined in (C) is divided into high bound MCM gate (>10^3^) which includes cells retaining high MCM intensity, and low bound MCM gate (<10^3^) which includes cells that have already unloaded MCM.

**Supplementary fig. 2.**
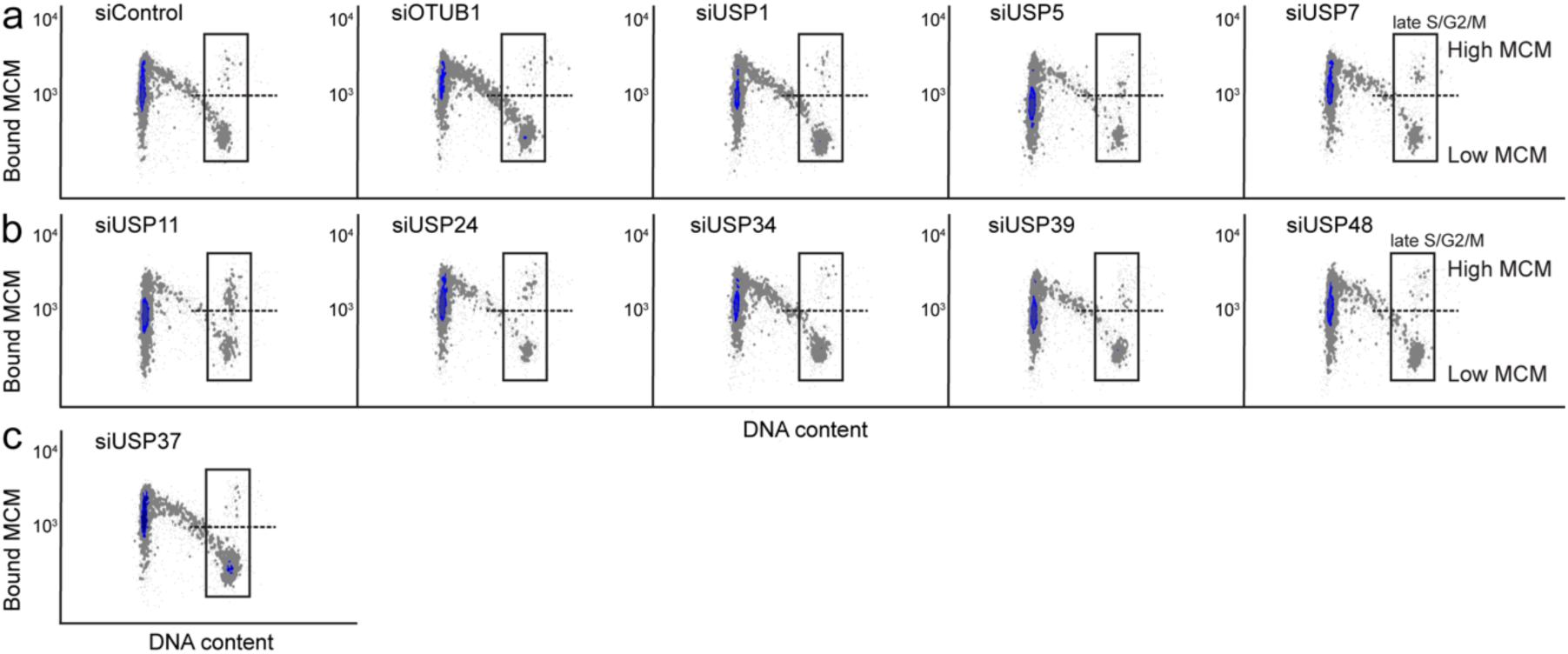
Bound MCM versus DNA content flow cytometry plots for the siRNA DUB screen. **A-C.** Chromatin flow cytometry for RPE1-hTert cells treated with 20 nM siControl or a panel of siRNAs targeting selected DUBs for 24 hours. Cells were stained for bound MCM2, and DAPI (for DNA content). Cells in each sample were randomly down sampled to 4000 cells per sample. All plots are from the same flow cytometry run. Representative of two biological replicates.

**Supplementary fig. 3.**
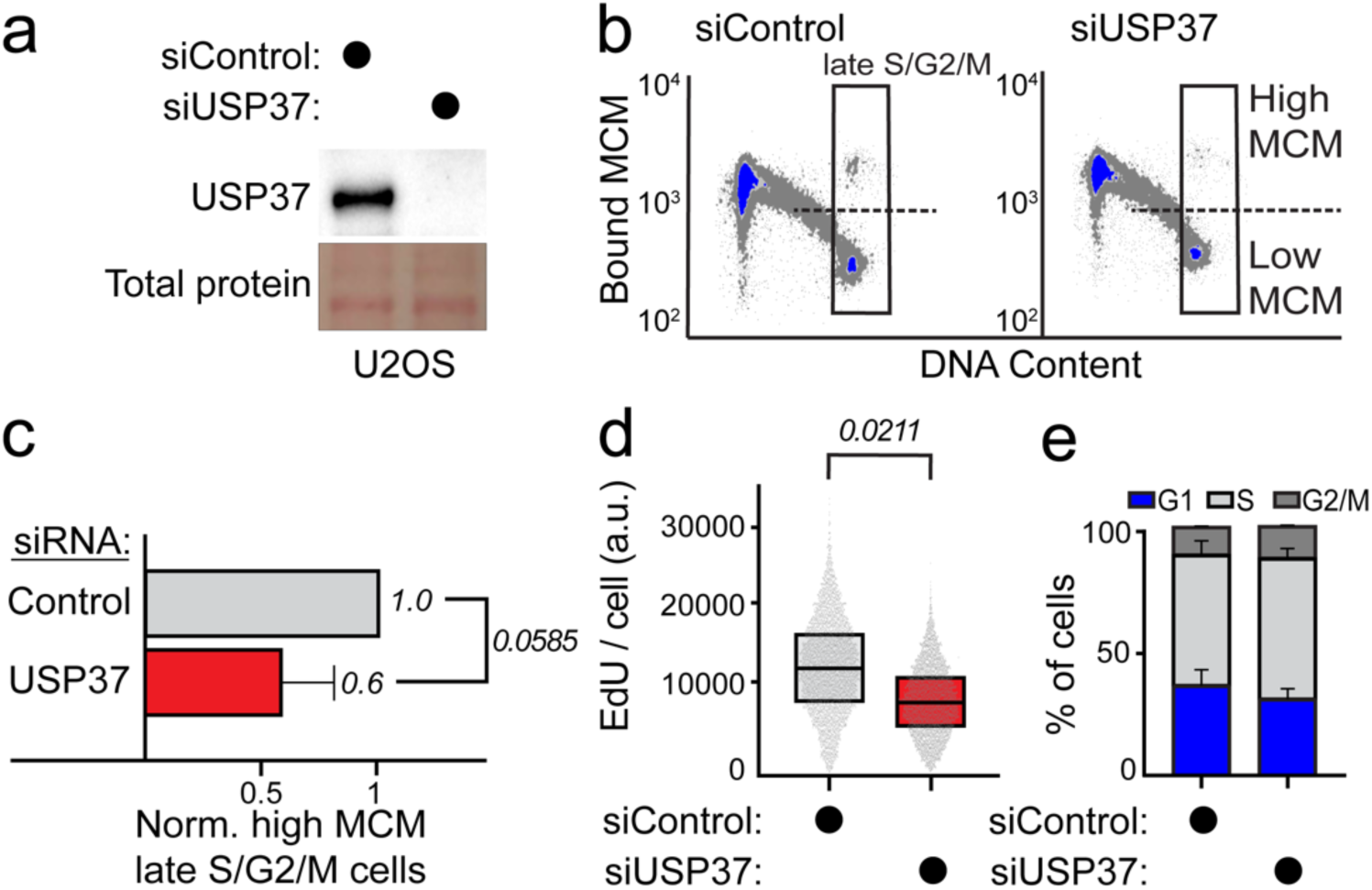
USP37 preserves loaded MCM in late S/G2/M and ensures replication progression in U2OS cells. **A.** Immunoblotting for USP37 in U2OS cells treated with 20 nM siControl or siUSP37 for 24 hours. Representative of two biological replicates. **B.** Chromatin flow cytometry for the same cells as in (A). Cells were pulsed with EdU for 30 min before harvesting. Cells were stained for bound MCM2, EdU incorporation (for DNA synthesis) and DAPI (for DNA content). In the late S/G2/M gate, control cells show more cells in the high chromatin bound MCM gate while cells depleted for USP37 show less cells in the high chromatin MCM gate. Cells were randomly down sampled to 15,000 cells per sample. Representative of two biological replicates. **C.** Relative percentage of high MCM, late S/G2/M-MCM^DNA^-positive cells from two biological replicates; mean with error bars ± SEM, unpaired two tailed t-test, p=0.0585. **D.** Box and whisker plots for EdU intensity per cell in S phase from the same samples as in (C). The aggregate of two biological replicates was randomly down sampled to 4000 cells per sample. Relative fold-change of the means of EdU intensity from the two replicates was computed, unpaired two tailed t-test, p=0.0211. **E.** Stacked bar graphs of the cell cycle phase distribution from the two biological replicates; mean with error bars ± SEM.

**Supplementary fig. 4.**
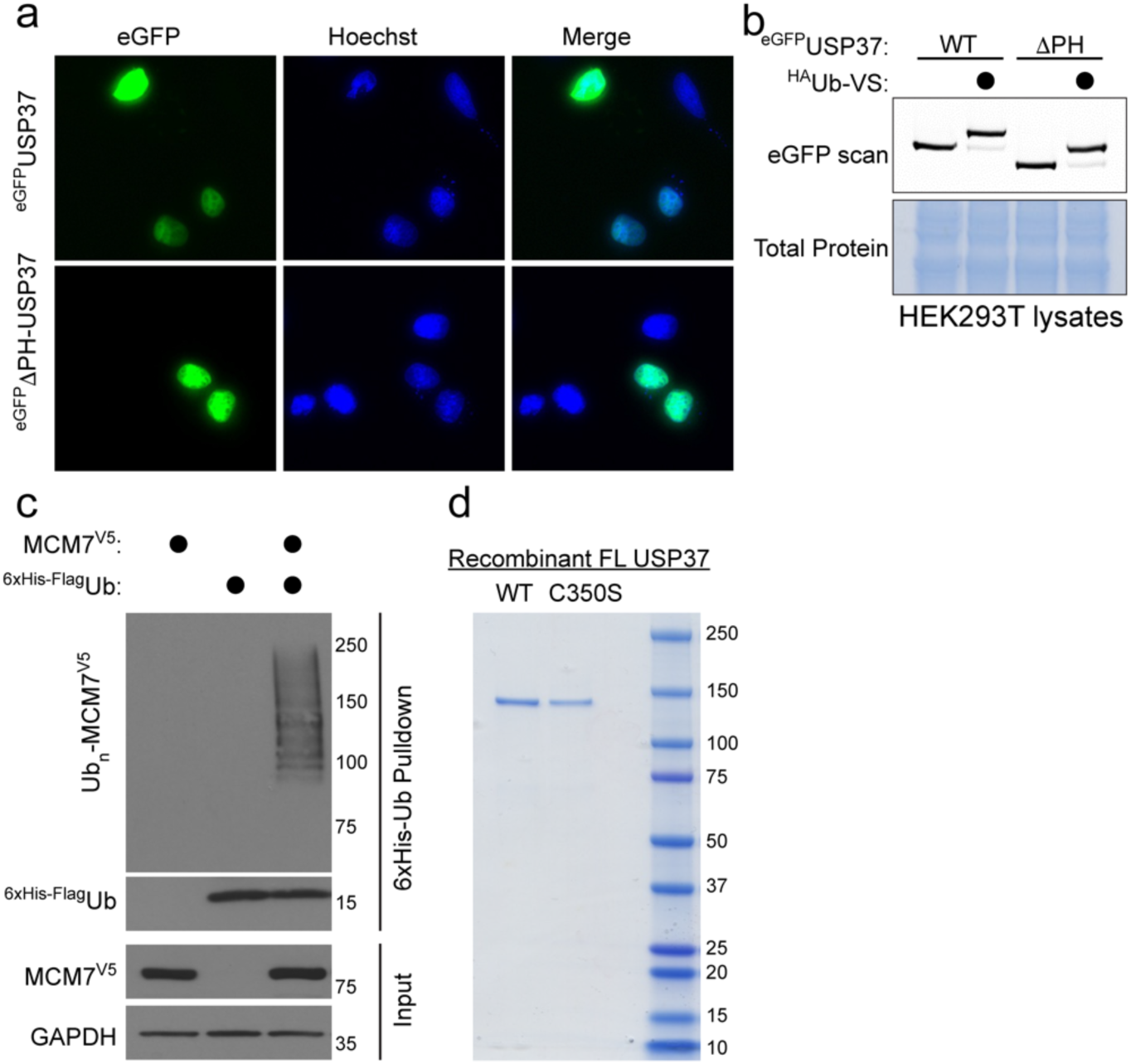
**A.** eGFP-tagged USP37 FL and Δ were ectopically expressed in U2OS cells for 24 hours. The day after, cells were seeded on glass coverslips and after 48 hours, cells were fixed and imaged using confocal microscopy. Both FL and Δ show similar nuclear localization. **B.** eGFP-tagged USP37 FL or Δ were ectopically expressed in HEK293T cells. After 24 hours, cell lysates were prepared and mixed with 5 µM of the ^HA^Ub-VS probe. Fluorescent scan shows that both USP37 FL and Δ have similar enzymatic capability. **C.** HEK293T cells were transiently transfected with a 6His-FLAG-tagged ubiquitin construct alone, a MCM7-V5 construct, either alone or in combination. Ubiquitinated proteins were isolated using Ni-NTA purification under strong denaturing conditions. MCM7 is heavily ubiquitinated under these conditions, as observed by immunoblotting. **D.** Recombinant USP37 WT or its catalytically inactive mutant (C350S) produced in insect cells were stained by Coomassie Brilliant Blue staining.

## MATERIAL AND METHODS

### Cell culture

All cells were tested for mycoplasma and confirmed negative. RPE1-hTERT, U2OS, and HEK293T cells were cultured and incubated in Dulbecco’s modified Eagle Medium (DMEM) with 10% fetal bovine serum (FBS), 2 mM L-glutamine, and 1x Pen-Strep at 37^°^C in a 5% CO_2_ incubator. RPE1-hTERT and U2OS cells were used for flow cytometry experiments. HEK293T cells were used for lentivirus packaging and transient transfections.

### Molecular biology

pDEST-HA-FLAG USP37 was a kind gift from Dr. Wade Harper (Addgene, cat. #22602). USP37 and mutants were subcloned into pcDNA3.1 (+) with an N-terminal Flag tag using Gibson assembly. To generate the EGFP-USP37 full length or ΔPH constructs, EGFP was first subcloned from a pCCL-EGFP vector (described in ^67^) into an empty pcDNA3.1(+) vector (Thermo Fisher, cat. #V79020) using HindIII and BamHI restriction sites to generate the pcDNA3.1(+)-EGFP vector. Then, USP37 and mutants were inserted using Gibson assembly. Doxycycline-inducible, siRNA resistant USP37^R^ was then made by two consecutive Gibson assembly reactions to introduce silent mutations corresponding to two siRNAs. First, the primer pair 5’- GCAACAGAACTCAGTCTTCAAGAGTTTAACAACTCCTTTGTGGATGCATTGG-3’ and 5’-CTCTTGAAGACTGAGTTCTGTTGCGCGCTTGAGGTCATCATCTTCTTTCTGTTCC- 3’ were used to alter coding starting at aa833. Simultaneously, the Flag tag was removed. In a second, separate reaction, the primer pair 5’- CCAGAGCCTATACATGTCCAGTGATTACTAATTTGGAGTTTGAGGTTCAGC-3’ and 5’-CCAAATTAGTAATCACTGGACATGTATAGGCTCTGGTAGCTGAAATATCTGG-3’ were used to alter the coding sequence starting at aa470. All constructs were confirmed by Sanger sequencing covering the full insert (Azentra). pLenti6-MCM7-V5 was a kind gift from Dr. Lynda Chin (Addgene, cat. #31212).

### Immunoblotting

After harvesting, cell pellets were washed with cold 1x PBS and lysed for 30 minutes in cold CSK buffer: (300 mM sodium chloride, 200 mM sucrose, 3 mM magnesium chloride, and 10 mM PIPES pH 7.0), which was supplemented with 0.5% triton X-100 (Sigma-Aldrich Chemistry) and a mixture of protease inhibitors: (1 µg/mL pepstatin A, 0.1 mM AEBSF, 1 µg/mL aprotinin and 1 µg/mL leupeptin), and phosphatase inhibitors: (1 mM β-glycerol phosphate, 10 µg/mL phosvitin and 1 mM Na-orthovanadate). Samples were cold centrifuged at maximum speed for 7.5 minutes. Protein content in the supernatants was quantified using Bradford assay (Biorad). Samples were diluted in SDS loading buffer to a final concentration of 1% SDS and 2.5% beta-mercaptoethanol, and boiled for 5 minutes. 20 ug of protein was run on 8% SDS-polyacrylamide gels, followed by wet transfer onto polyvinylidene difluoride (PVDF) membranes (Thermo Fisher Scientific). Ponceau S staining (Sigma-Aldrich) was used as the loading control in all experiments. Membranes were then blocked for 1 hour at room temperature in 5% milk diluted in Tris-buffered-saline-0.1% Tween 20 (TBST). Primary antibody was diluted in 2.5% milk-TBST and incubated with membranes overnight at 4°C. Blots were washed the next day 3x with 1x TBST and incubated with the secondary antibody for 1 hour at room temperature. Blots were washed 3x with 1x TBST then incubated with ECL Prime (Amersham) for 5 minutes and imaged using a Chemidoc Imaging system (BioRad).

The following antibodies were used in this study: primary antibodies; *USP37* (1:2000, Bethyl laboratories, A300-927A), *Cyclin E1* (1:2000, Cell Signaling Technology, 4129), *c-MYC* (1:1000, Invitrogen, clone 9E10, MA1-980), *MCM7* (1:1000, Cell Signaling Technology, 3735), *CDC45* (1:1000, Cell Signaling Technology, 3673), *MCM2* (1:10,000, BD, 610700), *GINS1* (1:1000, EMD Millipore, MABE2033), *GINS2* (1:500, ABClonal, A9172), *Ubiquitin* (1:5000, Cell Signaling Technology, 3933S), TIMELESS (1:1,000, Santa Cruz, #sc-393122), V5 (1:5,000, Thermo, # R960-25) and secondary antibodies; donkey anti-rabbit IgG HRP-conjugated (1:10,000, Jackson ImmunoResearch, 711-035-152), donkey anti-mouse IgG HRP-conjugated (1:10,000, Jackson ImmunoResearch, 715-035-150).

### Quantitative chromatin-bound MCM and CDC45 flow cytometry

To label cells that are actively dividing, cells were incubated with 10 μM of EdU (Santa Cruz Biotechnology) for 30 minutes prior to harvesting. Cells were harvested and soluble proteins were pre-extracted on ice for 8 minutes using CSK buffer: (300 mM sodium chloride, 200 mM sucrose, 3 mM magnesium chloride, and 10 mM PIPES pH 7.0) supplemented with 0.5% triton X-100 and a mixture of protease and phosphatase inhibitors as discussed above. Cells were washed in 1% BSA-PBS and fixed in 4% paraformaldehyde (Sigma-Aldrich Chemistry) diluted in PBS for 15 minutes at room temperature. Cells were washed in 1% BSA-PBS and stored at 4°C until staining.

For EdU detection, cells were incubated in EdU labelling solution: (1 μM Alexa-fluor 647 or 488 azide (Life Technology), 1 mM CuSO_4_, and 100 mM ascorbic acid diluted in PBS) for 30 minutes at room temperature in the dark. Cells were washed in 1% BSA-PBS + 0.1% NP-40 solution. For primary antibody staining, cells were incubated in MCM2 antibody (1:190, BD biosciences, cat. #610700) or CDC45 antibody (1:50, Cell Signaling Technology, cat. #11881) diluted in 1% BSA-PBS + 0.1% NP-40 for 1 hour at 37°C in the dark. Cells were washed in 1% BSA-PBS + 0.1% NP-40 solution. For secondary antibody staining, cells were incubated in donkey anti-mouse secondary antibody conjugated to Alexa-fluor 488 for MCM2 or donkey anti-rabbit secondary antibody conjugated to Alexa-fluor 647 for CDC45 (1:1000, Life Technology) for 1 hour at 37°C in the dark. Cells were washed in 1% BSA-PBS + 0.1% NP-40 solution. Finally, cells were incubated in 1 μg/mL DAPI (Sigma-Aldrich Chemistry) and 100 μg/mL RNase (Sigma-Aldrich Chemistry) diluted in 1% BSA-PBS + 0.1% NP-40 for 1 hour at 37°C in the dark or overnight at 4°C. Data were collected the next day on Attune NxT Flow cytometer and analyzed with FCS Express 7 software. Data analysis was performed as described in^35^. Control samples were not incubated in EdU labelling solution or MCM2 or CDC45 antibody to determine thresholds for positive EdU and MCM gating (Supplementary Fig. #1).

#### Flow cytometry statistical analysis

For the MCM quantification, the percentage of cells in the high MCM gate in late S/G2/M was determined with FCS Express 7 software for each sample in each biological replicate. Control was set to 1 and the relative fold-change in the percentage of high MCM cells was computed. *For the CDC45 and EdU quantification*, GraphPad Prism v10 was used to calculate the means of the single-cell fluorescence intensities for each sample in each biological replicate. Relative fold-change among the means was computed for pairwise comparisons. Unpaired two-tailed t-test was used to calculate p-values, assuming equal standard deviation (without Welch’s correction). Statistics were applied only to the means of independent experiments and not to single cells within an experiment. Outliers were removed using ROUT (Q = 2%) when necessary.

### siRNA transfection

Appropriate siRNAs were transfected into cells using Lipofectamine RNAiMAX (Invitrogen) according to manufacturer’s instructions. Briefly, siRNAs or Lipofectamine were individually mixed in Opti-MEM (Gibco), and then added together as a mixture to target cells in antibiotic-free DMEM supplemented with 10% FBS and L-glutamine after aspirating the original culture media. Samples were collected 24 hours after siRNA treatment and/or doxycycline addition. For the siRNA screen in Fig. 1 and for Supplementary Fig. 3, a mixture of 4 siRNAs were used for USP37 knockdown (5 nM each). For the rescue experiments in Fig. 2 and for Fig. 5, 5 nM of siUSP37-1 was used. The siRNAs used in this study were synthesized by Thermo Scientific.

The siRNA sequences and their final concentrations are as follows: siLuciferase (control siRNA):

siLuciferase (control siRNA): 5 or 20 nM, 5’ UCGAAGUACUCAGCGUAAG 3’
*siUSP37-1*: 5 nM for the siRNA screen and for the rescue experiments: 5’ CAAAAGAGCUACCGAGUUA 3’
*siUSP37-2:* 5 nM, 5’ GCAUACACUUGCCCUGUUA 3’
*siUSP37-3:* 5 nM, 5’ AAACAAAGCCGCCUAAUGU 3’
*siUSP37-4: 5 nM,* 5’ GAGGAUCGAUUAAGACUGU
*siUSP1 pool:* 20 nM, 5’ GCAUAGAGAUGGACAGUAU 3’, 5’ GAAAUACACAGCCAAGUAA 3’, 5’ CAUAGUGGCAUUACAAUUA 3’, 5’ GCACAAAGCCAACUAACGA 3’
*siUSP5 pool:* 20 nM 5’ GAGCUGACGUGUACUCAUA 3’, 5’ GGACAACCCUGCUCGAAUC 3’, 5’ GGAGAGACAUUUCAAUAAG 3’, 5’ GAUCUACAAUGACCAGAAA 3’
*siUSP7 pool:* 20 nM, ,5’ CUAAGGACCCUGCAAAUUA 3’, 5’ GUGGUUACGUUAUCAAAUA 3’, 5’ UGACGUGUCUCUUGAUAAA 3’, 5’ 3’
*siUSP11 pool:* 20 nM 5’ GGGCAAAUCUCACACUGUU 3’, 5’ GAACAAGGUUGGCCAUUUU 3’, 5’ GAUGAUAUCUUCGUCUAUG 3’, 5’ GAGAAGCACUGGUAUAAGC 3’
*siUSP24 pool:* 20 nM, ,5’ GGACGAGAAUUGAUAAAGA 3’, 5’ AGGGAAACCUUACCUGUUA 3’, 5’ CCACAGCUUUGUUGAAUGA 3’, 5’ GUAGAAGCCUUGUUGUUCA 3’
*siUSP34 pool:* 20 nM, 5’ GAAAUUGACUCUCCUUAUU 3;, 5’ UAACAUGGCUGACUUAAUG 3’, 5’ GCAAUGAGGUUAAUUCUAG 3’, 5’ GGACCAAAUUUACAUAUUG 3’
*siUSP39 pool:* 20 nM, 5’ GAUCAUCGAUUCCUCAUUG 3’, 5’ CAAGUUGCCUCCAUAUCUA 3’, 5’ UCACUGAGAAGGAAUAUAA 3’, 5’ ACAUAAAGGCCAAUGAUUA 3’
*siUSP48 pool:* 20 nM, 5’ CUACAUCGCCCACGUGAAA 3’, 5’ GCACUCUACUUAUGUCCAA 3’, 5’ GGCAGAGAGUCUAAGCUUU 3’, 5’ CGAAUUGCUUGGUUGGUAU 3’
*siOTUB1 pool:* 20 nM, 5’ GACGGACUGUCAAGGAGUU 3’, 5’ GACGGCAACUGUUUCUAUC 3’, 5’ CCGACUACCUUGUGGUCUA 3’, 5’ GACAACAUCUAUCAACAGA 3’

### Cell line generation

Lentiviral expression plasmids used were: pINDUCER20-USP37, pINDUCER20-cyclin E1, pINDUCER20-HA-HA-c-Myc, pCCL-WPS-mPGK_6his-FLAG-ubiquitin vectors^67^. To generate RPE1-hTERT cells stably expressing siRNA-resistant USP37 or cyclin E1 or HA-HA-c-Myc or 6his-FLAG-ubiquitin, lentivirus stocks were generated by co-transfecting HEK293T cells with the indicated lentiviral expression plasmid in addition to VSVG and ΔNRF (gifts from J. Bear) virus packaging plasmids using 50 μg/mL polyethylenimine (PEI)-Max (Sigma Aldrich Chemistry). RPE1-hTERT cells were transduced with the appropriate collected lentivirus using 8 μg/mL polybrene (Millipore) for 24 hours. Cells transduced with any of the pINDUCER20 plasmids were drug-selected using 500 μg/mL geneticin (Gibco) for 5 days. To pick individual clones from RPEs expressing siRNA-resistant USP37 or HA-HA-c-Myc, 2500 cells were plated sparsely in a 15 cm dish for clonal selection. Protein expression was confirmed by immunoblotting.

### Immunoprecipitation

#### Exogenous IP

For immunoprecipitation experiments using overexpressed proteins, DNA constructs were transfected using PolyJet (SignaGen) in HEK293T cells that were seeded in 10 cm dishes. After 48h, cells were washed in PBS, harvested in PBS, then pelleted by centrifugation at 1,500×g for 3 minutes at 4°C. Cell pellets were lysed on ice for 10 minutes using NETN lysis buffer supplemented with 10 μg/mL aprotinin, 10 μg/mL leupeptin, 10 μg/mL pepstatin A, 1 mM sodium orthovanadate, 1 mM sodium fluoride, and 1 mM AEBSF (4-[two aminoethyl] benzenesulfonyl fluoride). Lysates were clarified by centrifugation at maximum speed (14,000 rpm) for 10 minutes at 4°C using a benchtop microcentrifuge, and protein concentration was determined using Bradford (Biorad). Prior to IP, 10% of the total protein was removed as the input, while the remaining lysate was mixed with 25 μL of Anti-FLAG M2 Affinity Gel (Sigma, cat. #F2426) to isolate FLAG-tagged USP37. Immunoprecipitations were performed for 2h at 4°C while rotating, after which the beads were washed 4 times using NETN lysis buffer. After the final wash, the beads were resuspended in 2x Laemmli sample buffer and boiled at 95°C for 5 minutes. Input and IP samples were separated by SDS-PAGE and analyzed by immunoblotting.

#### USP37 IP-MS

To define the interactome of USP37, FLAG-EV or FLAG-USP37 was transfected into HEK293T cells seeded in 10 cm dishes, in triplicate, using 2 plates per condition. Cells were transfected with 2.5 µg of FLAG-USP37 using PolyJet (SignaGen) according to the manufacturer’s instructions. The day after, cells were amplified by transferring them to 15 cm dishes. After 48h of expression, cells were washed with PBS, collected and lysed in NETN lysis buffer supplemented with 10 μg/mL aprotinin, 10 μg/mL leupeptin, 10 μg/mL pepstatin A, 1 mM sodium orthovanadate, 1 mM sodium fluoride, and 1 mM AEBSF (4-[two aminoethyl] benzenesulfonyl fluoride). Lysates were snap frozen 2x using liquid nitrogen and clarified by centrifugation at 14,000 rpm for 15 minutes at 4°C. Protein concentration was determined and normalized using Bradford assay, and 18 mg of protein per sample were used. Samples were mixed with 50 µl of EZview Anti-FLAG M2 Affinity Gel (Sigma, cat. #F2426), and immunoprecipitation was performed for 4h at 4°C. After IP, samples were washed 3x using NETN lysis buffer followed by 3 washes using PBS. Beads were covered in PBS and frozen at –80C until further analysis by mass spectrometry (see below).

### Mass spectrometry

Immunoprecipitated protein samples were subjected to on-bead trypsin digestion as previously described^68^. Briefly, after the last wash step of the immunoprecipitation, beads were resuspended in 50µl of 50mM ammonium bicarbonate, pH 8. On-bead digestion was performed by adding 1µg trypsin and incubated with shaking, overnight at 37°C. The following day, 1ug trypsin was added to each sample and incubated shaking, at 37°C for 3 hours. Beads were pelleted and supernatants were transferred to fresh tubes. The beads were washed twice with 100µl LC-MS grade water, and washes were added to the original supernatants. Samples were acidified by adding TFA to final concentration of 2%, to pH ∼2. Peptides were desalted using peptide desalting spin columns (Thermo Scientific), lyophilized, and stored at -80°C until further analysis.

#### LC-MS/MS

Immunoprecipitation samples were analyzed by LC-MS/MS using an Easy nLC 1000 coupled to a QExactive HF mass spectrometer (Thermo Scientific). Samples were injected onto an Easy Spray PepMap C18 column (75 μm id × 25 cm, 2 μm particle size) (Thermo Scientific) and separated over a 2 hr method. The gradient for separation consisted of 5–45% mobile phase B at a 250 nl/min flow rate, where mobile phase A was 0.1% formic acid in water and mobile phase B consisted of 0.1% formic acid in ACN. The QExactive HF was operated in data-dependent mode where the 15 most intense precursors were selected for subsequent fragmentation. Resolution for the precursor scan (m/z 300–1600) was set to 120,000, while MS/MS scans resolution was set to 15,000. The normalized collision energy was set to 27% for HCD. Peptide match was set to preferred, and precursors with unknown charge or a charge state of 1 and ≥ 7 were excluded.

#### Data analysis

Raw data files were searched against the reviewed human database (containing 20,396 entries), appended with a contaminants database, using Andromeda within MaxQuant (v1.6.15.0). Enzyme specificity was set to trypsin, up to two missed cleavage sites were allowed, and methionine oxidation and N-terminus acetylation were set as variable modifications. A 1% FDR was used to filter all data. Match between runs was enabled (5 min match time window, 20 min alignment window), and a minimum of two unique peptides was required for label-free quantitation using the LFQ intensities.

Perseus was used for further processing^69^. Only proteins with >1 unique+razor peptide were used for LFQ analysis. Proteins with 50% missing values were removed and missing values were imputed from normal distribution within Perseus. Log2 fold change (FC) ratios were calculated using the averaged Log2 LFQ intensities of ^Flag^USP37 IP compared to control IP, and students t-test performed for each pairwise comparison, with p-values calculated. Proteins with significant p-values (<0.05) and Log2 FC >1 were considered biological interactors.

Gene ontology analysis was performed on the top 5% of interactors determine by expression over control using Metascape^70^.

### In vivo ubiquitination assay

For experiments using RPE1, parental or cells stably expressing a 6His-Flag-Ubiquitin construct (RPE1-6HF-Ub) were seeded in 10-cm dishes and transfected with 20 nM of the indicated siRNAs the day after. Knockdown was performed for 48 hours and 5× 10-cm dishes of cells at ∼90% confluence were used for each experimental condition. For the experiment described in Fig 4B,C, cells were treated with 5 µM of the p97 inhibitor CB-5083 (Selleck Chem Cat. #S8101) for the final 4 hours prior to harvesting. The in vivo ubiquitination assay was then performed essentially as described previously with minor adjustments^71^. Briefly, cells were washed twice with 5 ml of PBS per dish and collected in 5 ml of PBS per experimental condition. 10% of the cell suspension (500 μl) was removed to prepare inputs using a standard cell lysis buffer containing 1% Tween20 supplemented with protease inhibitors. The remaining 90% was centrifuged at 1,000g for 5 mn and resuspended in 8 ml of 6M guanidine-HCl containing buffer supplemented with 10 mM β-Mercaptoethanol and 15 mM Imidazole. His_6_-tagged ubiquitinated proteins were then captured on Ni^2+^-NTA agarose beads (Qiagen, cat. #30210) overnight at 4 degrees. After extensive washes of the beads using 8M Urea containing buffers, pull-down eluates as well as inputs were separated on SDS-PAGE gels and analyzed by immunoblot. For experiments using HEK-293T, cells were seeded in 10 cm dishes and transfected as indicated using PolyJet (SignaGen) and following the manufacturer’s instructions. The in vivo ubiquitination assay was then performed exactly as described previously in ^72^.

### In vitro deubiquitination of Ub-MCM7 by recombinant USP37

#### Purification of recombinant USP37

Purification of FL USP37 (1-979) from baculoviral infected insect cells was performed as described in ^73^. In brief, USP37 was inserted into pFastbac vector containing an N-terminal GST tag, a TEV protease site, and a Flag tag, as well as a C-terminal 6xHis tag. Baculoviral infected *Tni* cells (Expression Systems) were harvested approximately 72 hours after infection and resuspended in a buffer containing 50mM Tris pH 7.6, 200mM NaCl, 2.5% glycerol, 5mM DTT, and protease inhibitors. After lysis by sonication, lysates were clarified by centrifugation for 1 hour at 35000 *x g*. Lysates were incubated in batch with GS4B resin (Genesee Scientific) for 1 hour at 4°C before elution with lysis buffer supplemented with additional 50 mM Tris pH 7.6 and 10mM glutathione. Isolated protein was subjected to GST tag removal overnight with treatment by TEV protease. USP37 was then further purified by anion exchange chromatography and finished with size-exclusion chromatography over an SD200 10/300 Increase (Cytiva) into a buffer containing 20 mM HEPES pH 8.0, 200 mM NaCl, 1 mM DTT.

#### Deubiquitination assay

For the assays described in Fig 4D, we first generated ubiquitinated MCM7 by transfecting 6× 10 cm of HEK-293T cells, with 2.5 µg of 6His-Flag-Ubiquitin and 2.5 µg of MCM7-V5 plasmids per plate using PolyJet (SignaGen) and following the manufacturer’s instructions. After 48 hours of transfection, cells were washed with PBS and harvested by scraping followed by centrifugation at 1,500 rpm for 3 min. Cell pellets corresponding to 3× 10 cm were resuspended in 12 ml of 6 M guanidine-HCl containing buffer supplemented with 10 mM β-Mercaptoethanol and 15 mM Imidazole. After sonication and filtering of lysates on 0.40 µm cell strainer (REF), His_6_-tagged ubiquitinated proteins were isolated on Ni^2+^-NTA agarose beads (Qiagen #30210) for 4 hours at RT. Beads were washed extensively using 8 M Urea containing buffers (see above), and after the last wash, beads were resuspended in 50 mM Tris pH 8.0, 50 mM NaCl, plus 0.1% Triton X-100. After 10 min of equilibration, beads were washed once with buffer without detergent, once with buffer including 0.1% Trixton X-100, after which His_6_-tagged ubiquitinated proteins were eluted in 50 mM Tris pH 8.0, 50 mM NaCl containing 250 mM Imidazole. Elution was performed twice for 15 min at RT, and 600 µl of eluate was ultimately collected then kept at -80 or used immediately for deubiquitination assay. The deubiquitination assay was then conducted as follows. Recombinant DUBs were diluted in DUB buffer (50 mM Tris pH 8.0, 50 mM NaCl, 10 mM DTT) and incubated at RT for 10 min. Ubiquitinated proteins isolated from HEK-293T cells were also diluted in DUB buffer and incubated at RT for 10 min (usually, 5 µl of ubiquitinated proteins and 5 µl of DUB buffer per time point were used). Reactions were started by mixing the DUB with the ubiquitinated sample, placed at 30 degrees, and aliquots were taken at the indicated time points then quenched with 4X Laemmli buffer. Reaction products were boiled and separated on SDS-PAGE gels and analyzed by immunoblot.

### Cleavage assay of K11, K48 or K63 Tetra-ubiquitin chains by Flag-USP37

The assay was conducted largely as described in ^74^. HEK-293T cells were seeded in 10 cm dishes and transfected the day after using PolyJet (SignaGen) and following the manufacturer’s instructions. Two 10 cm dishes were transfected with 5 μg of Flag-USP37 per plate, and cells were harvested 24 hours after transfection. Cells were lysed for ∼10 min in phosphate lysis buffer (50 mM NaH_2_PO_4_, 150 mM NaCl, 1% Tween-20, 5% Glycerol, pH 8.0) supplemented with 2 μg/ml pepstatin, 1 mM AEBSF [4-(2 Aminoethyl) benzenesulfonyl fluoride], 1 mM Na_3_VO_4_ and 10 mM DTT. After centrifugating debris for 10 min at 14,000 rpm, anti-Flag M2 beads (F2426-1ML Millipore Sigma) were added to the lysate for 1 hour to immunoprecipitate Flag-USP37. Beads were washed once with lysis buffer, once with PBS then once with DUB buffer (50 mM Tris pH 7.5, 50 mM NaCl, 10 mM DTT) and split into 3 different tubes. The beads were centrifuged, resuspended in 25 μl of DUB buffer and incubated for 10 min at room temperature. In parallel, K11, K48 or K63 Tetra-ubiquitin chains (UC-45, UCB-210, UC-310, R&D Sytems) were prepared in DUB buffer at a final concentration of 2 μM then mixed with the USP37 immunoprecipitates. Aliquots of 10 μl were collected at the indicated time points, quenched with 5 μl of 4X laemmli buffer and separated by SDS-PAGE then analyzed by immunoblot.

### Fluorescence of EGFP-USP37 FL or ΔPH

U2OS cells were seeded in 6 cm dishes to reach 90% of confluency the day of transfection. Cells were then transfected with 2.5 µg of EGFP-USP37 FL or ΔPH domain using PolyJet (SignaGen) and following the manufacturer’s instructions. The day after, cells were plated on glass cover slips in 6-well plate format and incubated for another 24 hours. After 48 hours of transfection, cells were washed twice with PBS, fixed with 3.7% formaldehyde in PBS for 10 min at RT, washed with PBS twice, permeabilized with 0.2% Triton X-100 in PBS for 3 min, washed with PBS twice again, and nuclei were stained with 10 µg/ml of Hoechst in PBS for 5 min. Cells were washed with PBS twice, then in water two more times and cover slips were mounted on cover slides and imaged using a Revolve microscope system (ECHO).

### Charging of EGFP-USP37 FL or ΔPH with HA-Ub-VS in HEK-293T cell lysates

The assay was conducted largely as described in ^75^. HEK-293T cells were seeded in 10 cm dishes and transfected with 5 µg of EGFP-USP37 FL or ΔPH domain using PolyJet (SignaGen) and following the manufacturer’s instructions. Cells were lysed 24 hours after transfection in lysis buffer (50 mM Tris-HCl, pH 7.4, 150 mM NaCl, 1 mM EDTA, 1% Triton X-100) supplemented with 2 μg/ml pepstatin, 1 mM AEBSF [4-(2 Aminoethyl) benzenesulfonyl fluoride], 1 mM Na_3_VO_4_ and 10 mM DTT. Lysis was performed on ice for ∼10 min, debris were centrifuged for 10 min at 14,000 rpm and protein concentration was determined using Bradford reagent (Bio-Rad). To monitor the reactivity of either EGFP-USP37 FL or ΔPH with ubiquitin, 10 μl of each lysate containing 20 μg of total protein was combined with 10 μl of reaction buffer (50 mM Na_2_HPO_4_, 500 mM NaCl and 10 mM DTT, pH 7.9) containing 10 μM of HA-Ub-VS (R&D Systems, cat. #U-212). Reaction mixtures were incubated at 37°C for 2 hours and quenched by addition of 10 μl of 4X sample buffer. Samples were immediately separated by SDS-PAGE and the gel was scanned for EGFP fluorescence using a Typhoon FLA 9500. Equal protein loading was visualized by QC Colloidal Coomassie Blue staining (Bio-rad) following the manufacturer’s instructions.

### DepMap data

Expression corrected CERES gene correlation scores to USP37 knockout were downloaded from The Cancer Dependency Map v24Q2, accessed on June 13^th^, 2024. Data was plotted in GraphPad Prism v10.

### Cell fitness assays

RPE1 hTERT cells expressing doxycycline inducible Cyclin E1 or c-Myc were used for cell viability assays. Cells were plated in 6cm dishes and allowed to expand for 24 hours, after which media was changed for media containing either 100 ng/mL (Cyclin E1) or 25 ng/mL (c-MYC) of doxycycline. After 24 hours of induction, 4000 cells/well were plated into 24 well plates. Either siControl or siUSP37 was transfected using RNAiMax and OptiMEM either with or without indicated doxycycline concentration for 24 hours. After transfection, media was replaced with fresh media with or without doxycycline for an additional 72 hours (Cyclin E1) or 24 hours (c-MYC). Cell viability/fitness/proliferation was measured with resazurin sodium salt (Sigma Aldrich, cat. #R7017, 44 µM final concentration), which was added for 2 hours before reading. Fluorescence intensity was measured with 570 emission and 590 excitation wavelengths. Data was background subtracted with wells containing media plus reagent but no cells. Each well was normalized to the averaged siControl wells without doxycycline. Values represent means of 3 biological replicates ± SEM. Significance was determined with one-way ANOVA.

### Chemical reagents/inhibitors

The following chemicals/inhibitors were used in this study: Doxycycline (dox) (CalBiochem, cat. # 32485) was used at 20 ng/mL for the rescue experiments, and 100 or 25 ng/mL for the Cyclin E1 or c-MYC overproduction experiments; p97 inhibitor (Selleck, cat. #S8101) was used at 1.25 μM for the flow cytometry experiments.

